# KLRE1 shapes antifungal immunity by tuning dendritic cell and T cell activation through distinct heterodimeric partners

**DOI:** 10.64898/2025.12.03.690303

**Authors:** Fabián Salazar, Ivy M. Dambuza, Emily A. Sey, Cecilia Rodrigues, Mariano Malamud, Jamie Harvey, Janet A. Willment, Salomé LeidundGut-Landmann, Gordon D. Brown

## Abstract

Dendritic cells (DCs) instruct adaptive immunity by integrating inflammatory cues to regulate co-signalling molecules and T cell activation. We recently discovered that the pattern recognition receptor Dectin-1 (Clec7a) controls expression of a network of killer lectin-like receptors (KLRs), including KLRI1 and KLRI2, which regulate adaptive immune responses. Here, we identify KLRE1 as a key regulator of DC phenotype and function, acting through distinct heterodimeric interactions with KLRI1 and KLRI2. KLRE1 is expressed by tissue-resident and bone marrow-derived DCs and is dynamically regulated by inflammatory signals, including fungal stimuli. Mechanistically, KLRE1:KLRI1 heterodimers promote DC activation, increasing CD40, CD80, CD86, and MHC-II, whereas KLRE1:KLRI2 heterodimers restrain activation. Differential regulation of KLRI1 and KLRI2 by inflammatory signals shapes heterodimer availability and immune outcomes under distinct conditions. Functionally, KLRE1 deficiency enhances DC activation and T cell responses *in vitro* and *in vivo* during *Candida albicans* infection. Strikingly, KLRI1- and KLRI2-deficient mice showed opposite survival outcomes during systemic candidiasis, despite similar fungal burdens, implicating KLRE1 heterodimers in disease tolerance rather than pathogen clearance. Protection in KLRI1-deficient mice was associated with a more balanced T cell response, whereas susceptibility to infection associated with KLRI2 deficiency resulted from increased T cell activation and migratory potential, leading to systemic inflammation and renal dysfunction. Thus, KLRE1 heterodimers fine-tune DC-driven T cell immunity, balancing protection and immunopathology during fungal infection.

## Main

Dendritic cells (DCs) are central orchestrators of adaptive immunity, acting as professional antigen-presenting cells that link innate recognition to T cell activation. Through the integration of three key signals, antigen presentation via peptide-MHC complexes, expression of co-stimulatory and co-inhibitory molecules, and secretion of T cell polarising cytokines, DCs fine-tune the type, quality and magnitude of T cell responses to infectious and non-infectious challenges^1^. Regulation of DC activation is essential to balance protective immunity with immune tolerance and to limit immunopathology during infection.

Co-signalling molecules such as CD40, CD80, and CD86 play a pivotal role in determining the outcome of T cell priming by shaping T cell expansion, differentiation, and effector function^2^. Regulation of expression of these molecules is controlled by a network of innate immune receptors, including pattern recognition receptors (PRRs) such as Toll-like receptors (TLRs) and C-type lectin receptors (CLRs), which detect pathogen-associated molecular patterns and initiate transcriptional programmes that drive DC activation^3^. While the roles and functions of many PRRs, including TLRs, in DC biology are well defined^4^, there is less known about how CLRs contribute to DC function and the regulation of adaptive immunity.

Dectin-1 (Clec7a), the prototypical member of the CLR family, was first characterised as a key mediator of antifungal immunity but is now recognised to play broader roles in host defence against diverse pathogens and in the pathogenesis of cancer, autoimmunity, neurological disorders, and developmental conditions^5-7^. We recently discovered that Clec7a regulates the expression of several CLRs in DCs, including a cluster of killer lectin-like receptors (KLRs) such as KLRI1 and KLRI2 (co-submitted manuscript^8^). KLRs are classically associated with natural killer (NK) cells, where they fine-tune cytotoxicity and cytokine production through paired activating and inhibitory receptors. Both KLRI1 and KLRI2 are expressed in NK cells, where they heterodimerise with KLRE1 to regulate cytotoxicity, however, their functions *in vivo* remain poorly understood^9,10^. Notably, we recently demonstrated that the KLRE1:KLRI1 heterodimer directly regulates CD4+ T cell responses (co-submitted manuscript^8^). However, the contribution of these KLR receptors to DC activation has not yet been explored.

In this study, we identify KLRE1 as a critical regulator of DC phenotype and function. We show that KLRE1 can exert opposing effects on DC activation depending on its heterodimeric partner. Engagement of KLRE1:KLRI1 heterodimers promotes co-stimulatory molecule expression, whereas KLRE1:KLRI2 heterodimers act as negative regulators of DC activation. Finally, using a systemic fungal infection model, we link these opposing regulatory mechanisms to functional outcomes *in vivo*. Together, our findings uncover KLRE1 as a novel checkpoint that fine-tunes DC activation and T cell immunity through distinct receptor partnerships, highlighting an additional layer of immune regulation with relevance for antimicrobial immunity and therapeutic modulation of immune responses.

## Results

### KLRE1 is expressed by dendritic cells and regulated by inflammatory signals

We have previously shown that KLRE1:KLRI1 heterodimers play a key role in regulating gastrointestinal immunity to fungal infection (co-submitted manuscipt^8^). To further explore the role of KLRE1 in immunity, we assessed KLRE1 protein expression in DCs from gut-draining lymph nodes by flow cytometry. Using both a KLRE1-specific antibody (validated on NIH3T3 cells overexpressing KLRE1 (co-submitted manuscript^8^), **Suppl. Fig. 1a**) and a KLRE1-EGFP knock-in reporter mouse (in which part of the KLRE1 genomic sequence was replaced with EGFP, resulting in a KLRE1 knockout)^11^, we detected significant KLRE1 protein expression in Lin^-^CD11c^+^MHC-II^+^ DCs (∼13–14% of cells) from mesenteric lymph nodes (mLNs) at steady-state (**Fig.1a** and **Suppl. Fig. 1b**). We also detected KLRE1 protein expression in splenic Lin^-^CD11c^+^MHC-II^+^ DCs under steady-state conditions, although at a lower frequency (∼3% of cells) (**Suppl. Fig. 1c**). In bone-marrow derived DCs we found that Flt3L drives the highest KLRE1 expression compared to standard cultures supplemented with GM-CSF, or GM-CSF combined with IL-4^12,13^ (**Fig. 1b**). Notably, culturing bone marrow cells on a stromal monolayer expressing the Notch ligand DLL1, that induces a tissue DC phenotype^14,15^, further increased KLRE1 expression (**Fig. 1b**). Interestingly, GM-CSF markedly suppressed KLRE1 expression; however, supplementation with Flt3L or IL-4 was sufficient to upregulate KLRE1 even in the presence of GM-CSF (**Suppl. Fig. 1d**). We confirmed KLRE1 protein expression in BMDCs under Flt3L+DLL1 conditions (FD-BMDCs) by flow cytometry (∼30% of cells) (**Fig. 1c**).

**Fig. 1.**
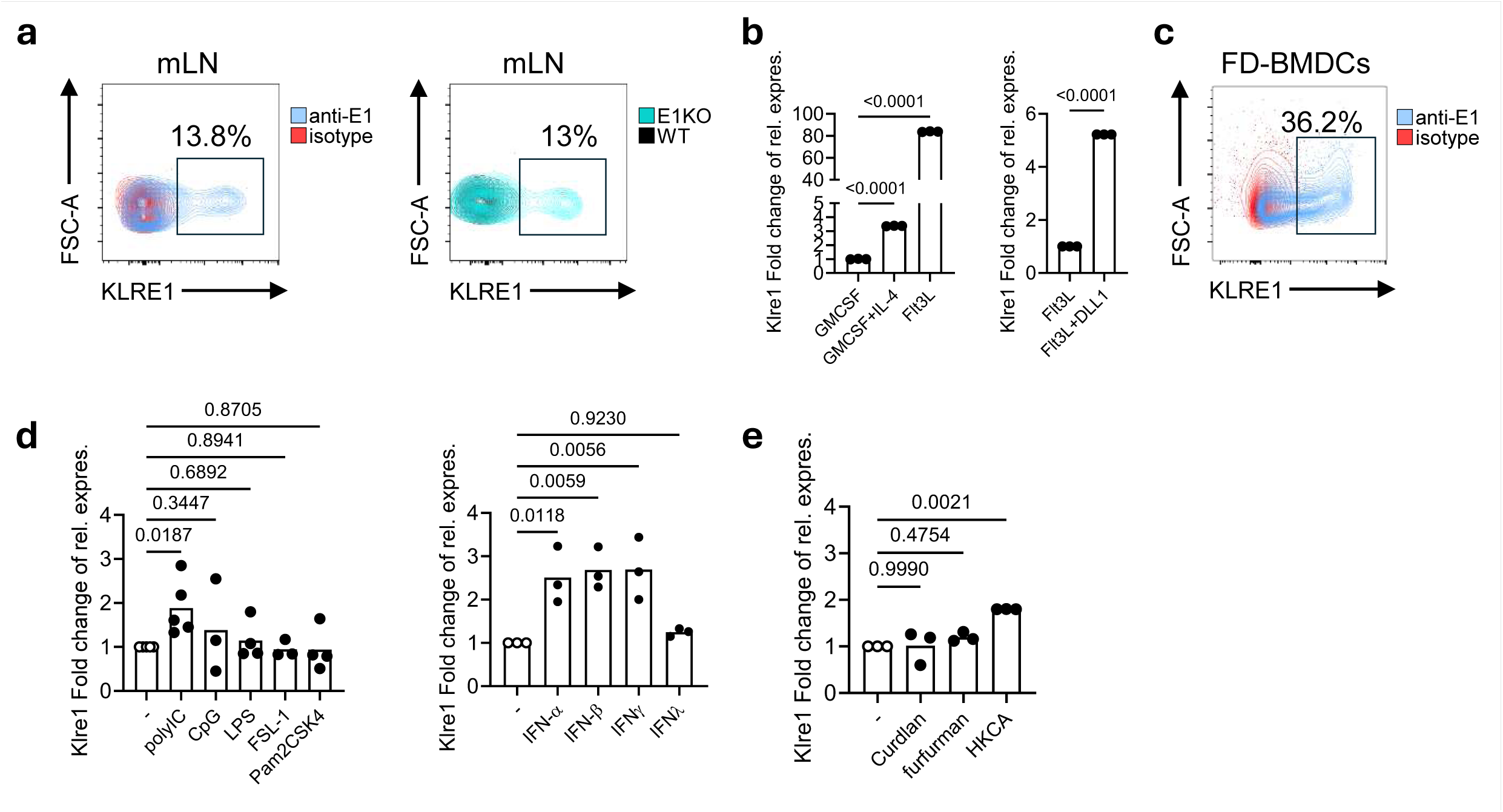
KLRE1 is expressed by DCs and regulated by inflammatory signals. **(a)** Flow cytometric analysis of KLRE1 expression in mLN DCs (gated on viable CD45+ Lin- CD11c+ MHC-II+ CX3CR1-). KLRE1 expression was assessed with a specific anti-KLRE1 monoclonal antibody (blue), using an isotype control for background (red), and compared to expression in KLRE1-EGFP knock-in mice (blue) with WT mice as controls (red). **(b)** KLRE1 gene expression in bone marrow–derived cells cultured for at least 7 days with different cytokine combinations. Data are shown as expression relative to GM-CSF (left) or Flt3L (right) treated samples and normalized to housekeeping genes GAPDH and 18srRNA. Bars represent the mean from one experiment with triplicates, shown as representative of at least two independent experiments. **(c)** Flow cytometric analysis of KLRE1 expression in BMDCs generated using the Flt3L-OP9 system (FD-BMDCs). KLRE1 expression was assessed with a specific anti-KLRE1 monoclonal antibody (blue), using an isotype control for background (red). **(d)** KLRE1 mRNA expression in FD-BMDCs after 16 h stimulation with indicated TLR ligands, or interferons. Data are shown relative to unstimulated controls and normalised to housekeeping genes. Bars represent mean of pooled data from at least three independent experiments. **(e)** KLRE1 mRNA expression in FD-BMDCs after 16 h stimulation with indicated CLR agonists. Data are shown relative to unstimulated controls and normalised to housekeeping genes. Bars represent mean of pooled data from at least three independent experiments. Statistical analysis was performed using Student’s t-test for two comparisons and One-way ANOVA for three or more comparisons; *p < 0.05.

We next investigated how various innate immune stimuli regulate KLRE1 expression. Among a panel of TLR ligands tested on FD-BMDCs, the TLR3 agonist polyinosinic:polycytidylic acid (polyI:C) significantly induced KLRE1 expression (**Fig. 1d**). where unmethylated CpG oligodeoxynucleotide (ODN), lipopolysaccharide (LPS), fibroblast-stimulating lipopeptide-1 (FSL-1), or S-[2,3-bis(palmitoyloxy)propyl]-Cys-Ser-Lys-Lys-Lys-Lys (Pam2CSK4) had little or no effect. Given that TLR3 signalling is a potent inducer of IFNs, we assessed how these cytokines would impact KLRE1 expression and found that both type I IFNs (IFN-α, IFN-β) and type II IFN (IFN-γ) robustly upregulated KLRE1 at the transcript and protein level (**Fig. 1d and Suppl. Fig. 1e**). We also examined the role of C-type lectin receptor (CLR) ligands in modulating KLRE1. Of the stimuli tested, only heat-killed *Candida albicans* (HKCA) induced KLRE1 transcription, while purified β -glucan- and mannan-containing molecules (e.g., curdlan and furfurman) had no effect (**Fig. 1e**). Together, these data demonstrate that KLRE1 expression in DCs is dynamically regulated by the inflammatory environment, particularly by signals involving TLR3, type I and II IFNs, and fungal pathogens.

### KLRE1 alters DC phenotype by regulating co-signalling molecule expression

To investigate the function of this CLR in DCs, we examined the effects of anti-KLRE1 capping on FD-BMDC activation. Using a TaqMan array, we found that antibody-mediated KLRE1 capping led to the transcriptional upregulation of multiple genes associated with DC activation. These included co-stimulatory molecules and additional components involved antigen presentation (*Cd40*, *Tapbp*, *Cd1d2*), cytokines, chemokines, and growth factors (*Il2*, *Ccl17*, *Cxcl2*, *Mdk*, *Il12b*), signalling molecules (*Relb*, *Stk4*, *Nfkb2*), as well as scavenger receptors and adhesion molecules (*Cd36*, *Icam1*) (**Fig. 2a**). Given the role of co-stimulatory molecules such as CD40 in shaping DC function and T cell activation, we specifically assessed the impact of KLRE1 capping on co-signalling molecule expression at the protein level. Antibody-mediated KLRE1 capping significantly upregulated surface expression of CD40, CD80, CD86, and MHC-II (**Fig. 2b** and **Suppl. Fig. 2a**), confirming that KLRE1 capping promotes a more activated DC phenotype. We next assessed the phenotype of KLRE1-deficient DCs with respect to co-signalling molecule expression. Strikingly, KLRE1-deficient FD-BMDCs showed significantly increased surface expression of CD80, CD86, and MHC-II, with CD40 showing an upward trend, compared to wild-type (WT) DCs (**Fig. 2c**), indicating that KLRE1 may act as a negative regulator of co-signalling molecule expression under resting conditions. Together, these results indicate that the loss or sequestration of KLRE1 through antibody capping, leads to a dysregulated enhanced expression of co-stimulatory molecules, establishing KLRE1 as a critical modulator of DC activation status.

**Fig. 2.**
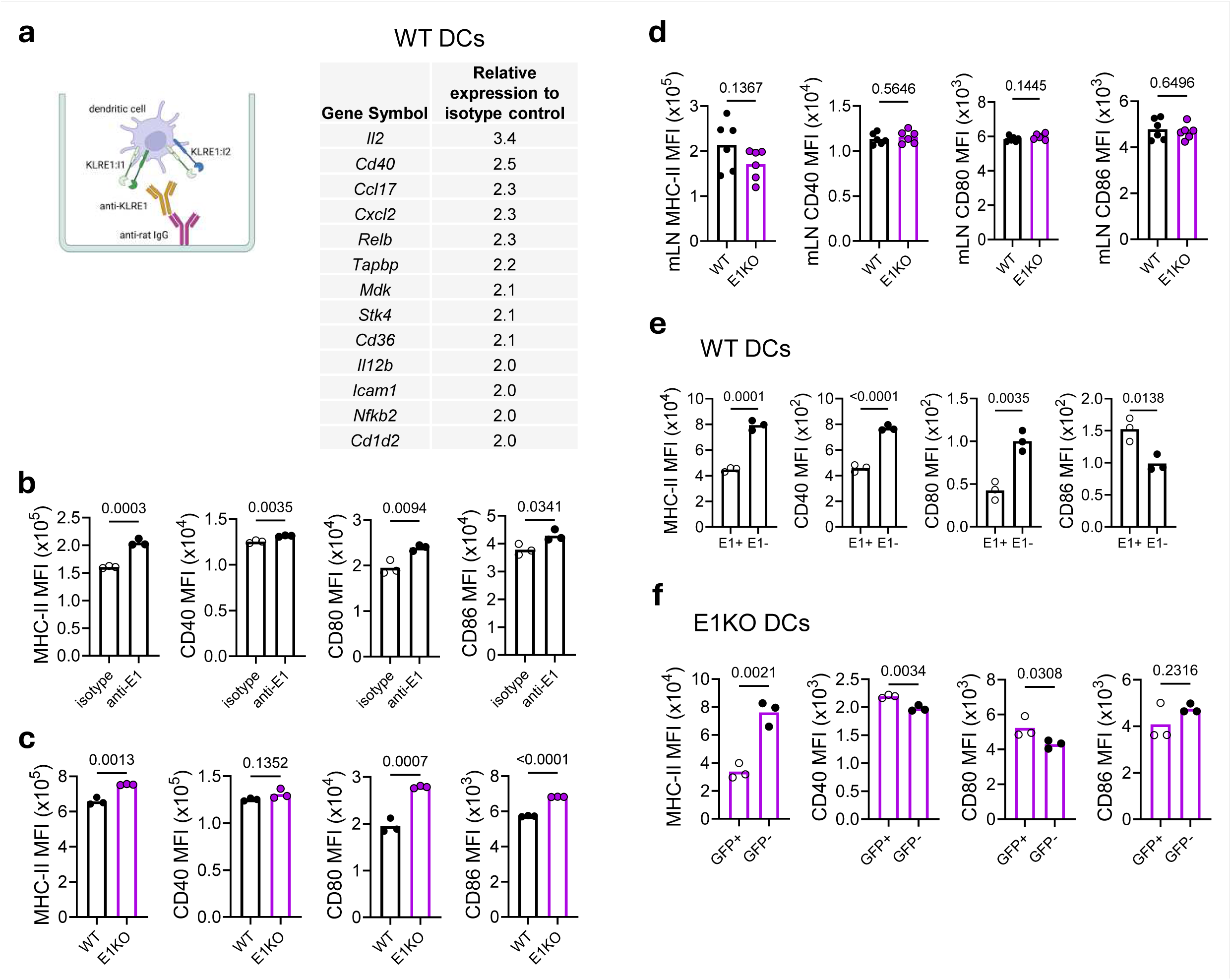
KLRE1 alters DC phenotype by regulating co-signalling molecule expression. (a) Transcriptional profiling of FD-BMDCs stimulated for 16 h with plate-bound anti-mouse KLRE1 antibody pre-capped with anti-rat IgG, compared with isotype control, using a TaqMan array. Table shows genes with ≥͑-fold upregulation. Data are shown as expression relative to isotype control and normalised to housekeeping genes. **(b)** Flow cytometric analysis of surface expression of MHC-II, CD40, CD80, and CD86 on FD-BMDCs after 24 h stimulation with plate-bound anti-mouse KLRE1 antibody pre-capped with anti-rat IgG. Data show median fluorescence intensity (MFI) from one experiment, shown as representative of at least two independent experiments. **(c)** Flow cytometric analysis of MHC-II, CD40, CD80, and CD86 expression in unstimulated FD-BMDCs from WT and KLRE1-deficient mice. Data show MFI from one experiment, shown as representative of at least two independent experiments. **(d)** Flow cytometric analysis of MHC-II, CD40, CD80, and CD86 expression in mLN DCs at steady state. Each point representsone mouse; horizontal bars indicate the mean. Data show MFI of pooled data from at least two independent experiments. **(e)** Flow cytometric analysis of MHC-II, CD40, CD80, and CD86 expression in KLRE1 expressing and KLRE1 non-expressing DC subsets isolated from mLN at steady state. Each point representsan individual mouse; horizontal bars indicate the mean. Data show MFI from one experiment, shown as representative of at least two independent experiments. **(f)** Flow cytometric analysis of MHC-II, CD40, CD80, and CD86 expression in mLN-derived GFP+ and GFP- DC subsets isolated from KLRE1-deficient mice at steady state. Each point representsan individual mouse; horizontal bars indicate the mean. Data show MFI from one experiment, shown as representative of at least two independent experiments. Statistical analysis was performed using Student’s t-test for two comparisons and Two-way ANOVA when assessing the effect of two independent variables; *p < 0.05.

We then examined whether this regulatory role extends to tissue-resident DCs. Analysing Lin^-^CD11c^+^MHC-II^+^CX3CR1^-^ DCs from naïve WT and KLRE1-deficient mice revealed no significant global differences in expression of CD40, CD80, CD86, or MHC-II, except for CD80 which was upregulated on splenic DCs (**Fig. 2d** and **Suppl. Fig. 2b-c**). Given that the level of expression of KLRE1 is limited to a small population of tissue DCs (**Fig. 1a**), which likely limits our ability to detect broad or global changes in co-signalling molecule expression, we next compared the expression of these molecules in KLRE1 expressing and non-expressing Lin^-^CD11c^+^MHC-II^+^CX3CR1^-^DCs from WT mice. We observed that WT DCs that do not express KLRE1 exhibited significantly higher levels of CD40, CD80, and MHC-II compared to DC that express this receptor (**Fig. 2e**). Importantly, these differences were largely lost (and even partially reversed) when comparing levels of expression of CD40, CD80 and CD86 in GFP+ vs GFP- Lin-CD11c+MHC-II+ DCs from the KLRE1-EGFP knock-out reporter mice (**Fig. 2f**). MHC-II levels remained high in GFP- compared to GFP^+^ Lin-CD11c+MHC-II+ DCs from the KLRE1-EGFP knock-out reporter mice. Taken together, these data support a model in which KLRE1 acts as a modulator of co-signalling molecule expression in DCs.

### KLRE1 modulates DC activation via distinct heterodimeric interactions with KLRI1 and KLRI2

KLRE1 has been shown to inversely regulate NK cell cytotoxicity *in vitro* by forming distinct heterodimers with receptors such as KLRI1 and KLRI2^10,11^. We have recently shown that KLRE1:KLRI1 heterodimers on DCs regulate T cell immunity during fungal infection (co-submitted manuscript^8^). Importantly, we found that KLRI1 and KLRI2 expression is highly dependent on KLRE1 as KLRE1-deficient mice show significant reduced expression of both KLRI1 and KLRI2 (**Suppl. Fig. 3a**). We next explored the possibility that the functions of KLRE1 in regulating DC phenotype might similarly depend on its pairing with different heterodimeric partners. To test this, we first examined whether KLRI1 and KLRI2 are coregulated with KLRE1 at the transcriptional level. We measured KLRI1 and KLRI2 gene expression in FD-BMDCs stimulated with various TLR and CLR agonists. Like KLRE1, both KLRI1 and KLRI2 were significantly upregulated by the TLR3 agonist polyI:C, but not by other TLR ligands such as CpG-ODN, LPS, FSL-1, or Pam2CSK4 (**Suppl. Fig. 3b-c**). Similarly, stimulation with HKCA significantly increased expression of both KLRI1 and KLRI2 (**Fig. 3a**). Notably, KLRI1 and KLRI2 showed differential regulation under other *in vitro* conditions: KLRI1, but not KLRI2, was robustly upregulated by IFN-y (**Suppl. Fig. 3b-c**). These data suggest that distinct inflammatory signals selectively regulate the availability of KLRE1 heterodimeric partners on DCs.

**Fig. 3.**
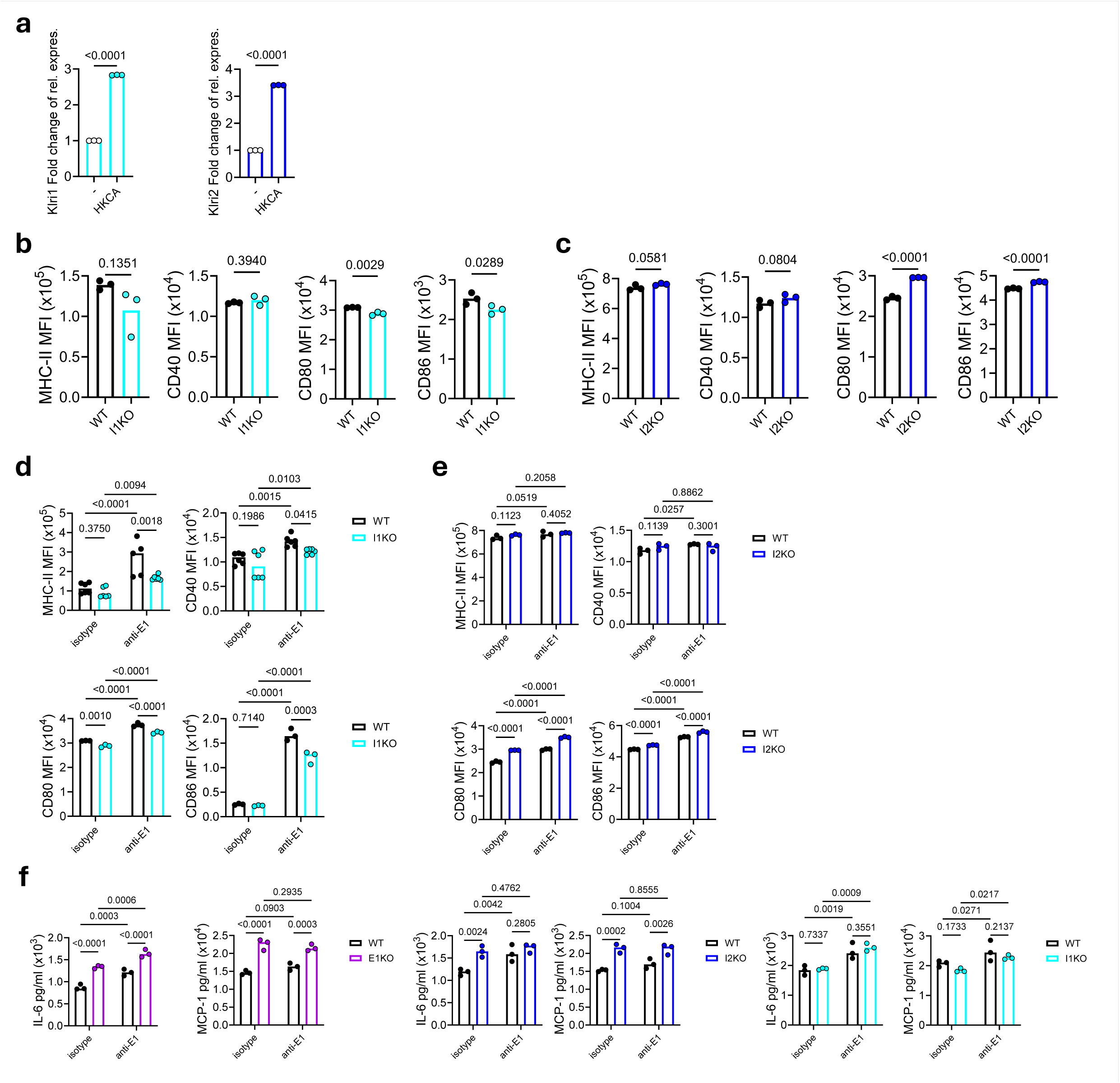
KLRE1 modulates DC activation via distinct heterodimeric interactions with KLRI1 and KLRI2. (a) KLRI1 and KLRI2 mRNA expression in FD-BMDCs after 16 h stimulation with HKCA. Data are shown relative to unstimulated controls and normalised to housekeeping genes. Bars represent the mean from one experiment, shown as representative of at least three independent experiments. **(b)** Flow cytometric analysis of MHC-II, CD40, CD80, and CD86 expression in unstimulated FD-BMDCs from WT and KLRI1-deficient mice. Data show MFI from one experiment, shown as representative of at least two independent experiments. **(c)** Flow cytometric analysis of MHC-II, CD40, CD80, and CD86 expression in unstimulated FD-BMDCs from WT and KLRI2-deficient mice. Data show MFI from one experiment, shown as representative of at least two independent experiments. **(d)** Flow cytometric analysis of MHC-II, CD40, CD80, and CD86 expression in FD-BMDCs from WT and KLRI1-deficient mice after 24 h stimulation with plate-bound anti-mouse KLRE1 antibody pre-capped with anti-rat IgG. Data show MFI from one experiment, shown as representative of at least two independent experiments. **(e)** Flow cytometric analysis of MHC-II, CD40, CD80 and CD86 expression in FD-BMDCs from WT and KLRI2-deficient mice after 24 h stimulation with plate-bound anti-mouse KLRE1 antibody pre-capped with anti-rat IgG. Data show MFI from one experiment, shown as representative of at least two independent experiments. **(f)** IL-6 and MCP-1 production by FD-BMDCs from WT, KLRE1-, KLRI2- and KLRI1-deficient mice after 24 h stimulation with plate-bound anti-mouse KLRE1 antibody pre-capped with anti-rat IgG. Horizontal bars indicate the mean from one experiment. Statistical analysis was performed using Student’s t-test for two comparisons and Two-way ANOVA when assessing the effect of two independent variables; *p < 0.05.

We next investigated how KLRE1 heterodimerisation with KLRI1 or KLRI2 influences DC phenotype. As previously shown (co-submitted manuscript^8^), loss of KLRI1 does not alter expression of KLRI2 or KLRE1, and KLRI2 deficiency likewise does not affect KLRI1 or KLRE1 expression (**Suppl. Fig. 3d**). We analysed co-signalling molecule expression in unstimulated FD-BMDCs from KLRI1- and KLRI2-deficient mice. While KLRI1-deficient cells showed reduced CD80, CD86 and MHCII expression, KLRI2-deficient FD-BMDCs closely resembled KLRE1-deficient FD-BMDCs, with increased expression of CD80, CD86, CD40 and MHC-II (**Fig. 3b-c**). Together, these findings indicate that KLRE1 forms distinct heterodimers with KLRI1 and KLRI2 that exert opposing effects as positive and negative regulators of DC co-stimulatory molecule expression, respectively.

To investigate this further, we re-examined the effect of KLRE1 capping in the absence of each heterodimeric partner. Based on the expression patterns of co-signalling molecules shown above, we anticipated reduced activation in KLRI1-deficient DCs and enhanced activation in KLRI2-deficient DCs compared to WT DCs. Indeed, KLRE1 capping in KLRI1-deficient DCs induced much less co-signalling molecule expression than in WT DCs, demonstrating that KLRE1:KLRI1 heterodimers positively contributes to KLRE1-mediated activation (**Fig. 3d**). In contrast, there was significantly higher levels of co-signalling molecule expression, particularly CD80 and CD86, following KLRE1 capping in KLRI2-deficient DCs (**Fig. 3e)**. Notably, we found that this dual regulation extended to cytokine production. KLRE1- and KLRI2-deficient DCs secreted significantly more IL-6 and MCP-1 than WT controls under steady-state conditions, and increased IL-6 production following KLRE1 capping (**Fig. 3f**). In contrast, KLRI1-deficient DCs produced cytokine levels similar to WT (**Fig. 3f**). Together, these results reveal that KLRE1 forms distinct heterodimers with KLRI1 or KLRI2, exerting opposing effects on DC activation.

### KLRE1 regulation of DC phenotype impacts T cell activation during fungal infection

We next investigated the functional consequences of KLRE1-deficiency on the ability of DCs to activate T cells. *In vitro*, co-culture experiments revealed that KLRE1-deficient DCs markedly enhanced OT-II T cell activation in response to OVA antigen, as shown by increased CD25 and CD44 expression (**Fig. 4a**). Similarly increased T cell activation was also observed following antibody-mediated capping of KLRE1 on WT DCs (**Suppl. Fig. 4a**).

**Fig. 4.**
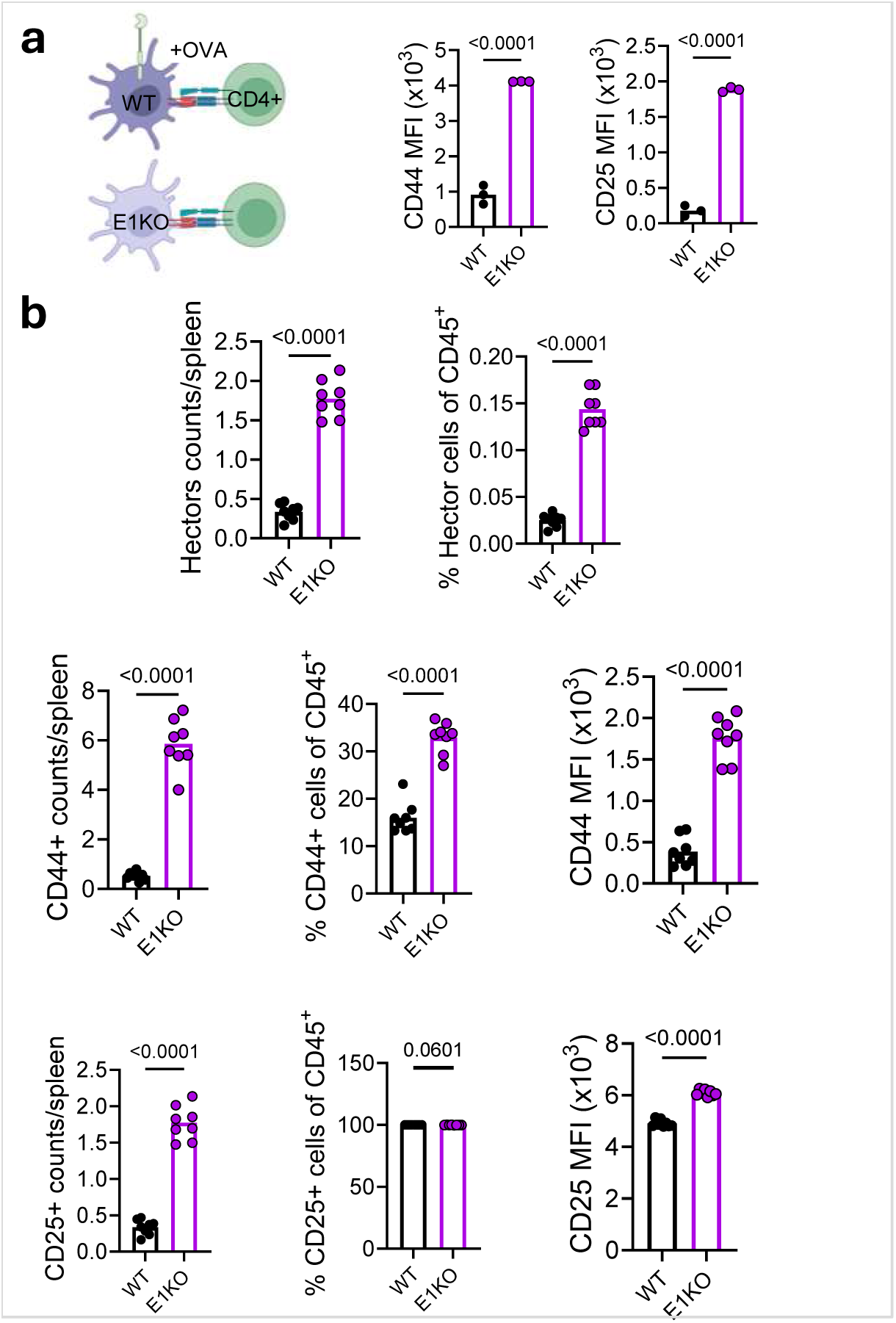
KLRE1 regulation of DC phenotype impacts T cell activation during fungal infection. (a) *In vitro* co-culture of WT or KLRE1-deficient DCs preloaded with OVA and OT-II CD4+ T cells for 72 h. T cell activation was assessed by CD44 and CD25 expression. Data show MFI from one experiment, shown as representative of at least two independent experiments. **(b)** Flow cytometric analysis of CD4+ T cells in spleen at day 3 post-infection, showing cell numbers, frequencies, and activation status based on CD44 and CD25 expression. Each point represents one mouse; horizontal bars indicate the mean. Data show MFI of pooled data from at least two independent experiments. Statistical analysis was performed using Student’s t-test for two comparisons and One-way ANOVA for three or more comparisons; *p < 0.05.

We then assessed the role of KLRE1 in the development of T cell responses *in vivo,* using adoptive transfer of *C. albicans*-specific Hector TCR transgenic T cells^16^ into WT and KLRE1-deficient mice to monitor antigen specific responses. KLRE1-deficient mice show no overt differences in immune cell populations within the mLN or bone marrow (**Suppl. Fig. 4b-c** and **Suppl. Fig. 4d-e**). We observed that following systemic infection with *Candida albicans*, KLRE1-deficient mice exhibited a pronounced increase in splenic antigen-specific T cell numbers, frequency, and activation, as indicated by elevated CD44 and CD25 expression compared to WT controls (**Fig. 4b** and **Suppl. Fig. 4f**). Collectively, these results demonstrate that KLRE1 plays a fundamental role in shaping T cell responses, acting as a regulator of immune activation.

### KLRI1 and KLRI2 inversely influence susceptibility to systemic *C. albicans* infection

To determine whether the increased T cell activation in KLRE1-deficient mice translates into improved protection against infection, we systemically infected WT and KLRE1-deficient mice and monitored progression of disease. Surprisingly, despite enhanced T cell activation (**Fig 4b**), KLRE1-deficient mice did not show altered survival compared to WT controls (**Fig. 5a**). Given that KLRE1 deficiency affects the expression of both KLRI1 and KLRI2, which we have shown to have opposing effects on DC activation, we hypothesised that their loss might counterbalance each other’s roles *in vivo*, masking any impact on susceptibility. To investigate this, we systemically infected KLRI1- and KLRI2-deficient mice with *C. albicans*. Strikingly, KLRI1-deficient mice showed significantly increased resistance to infection, while KLRI2-deficient mice were more susceptible (**Fig. 5b-c**).

**Fig. 5.**
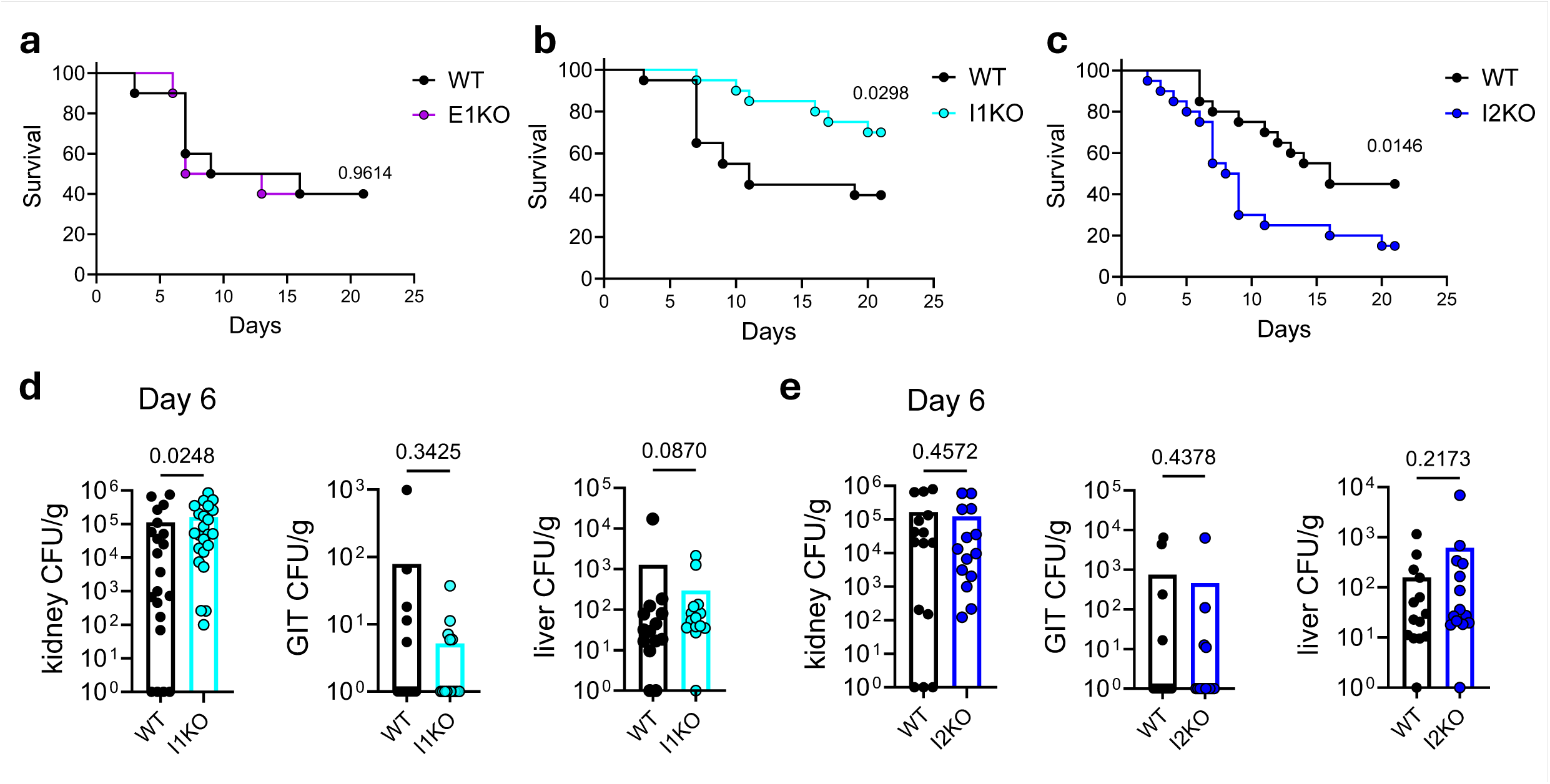
KLRI1 and KLRI2 inversely influence susceptibility to systemic *Candida* infection. (a) Survival of WT and KLRE1-deficient mice up to day 21 following systemic infection with 1 × 10⁵ CFU *C. albicans*. Data are from one experiment with ten mice per group. **(b)** Survival of WT and KLRI1-deficient mice up to day 21 following systemic infection with 1 × 10⁵ CFU *C. albicans*. Data are pooled from two independent experiments, with ten mice per group in each experiment. **(c)** Survival of WT and KLRI2-deficient mice up to day 21 following systemic infection with 1 × 10⁵ CFU *C. albicans*. Data are pooled from two independent experiments, with ten mice per group in each experiment. **(d)** Fungal burden (colony-forming units, CFUs) of kidney, GIT, and liver of WT and KLRI1-deficient mice at day 6 post-infection. Each point representsan individual mouse; bars indicate mean. Data pooled from at least two independent experiments, with ten mice per group in each experiment. **(e)** Fungal burden of kidney, GIT, and liver of WT and KLRI2-deficient mice at day 6 post-infection. Each point representsan individual mouse; bars indicate mean. Data pooled from two independent experiments, with ten mice per group in each experiment. Statistical analysis was performed using Student’s t-test for two comparisons. Fungal burden data were evaluated with Mann–Whitney U tests. Survival data were analysed with Kaplan-Meier test. *p < 0.05.

To determine whether these differences in survival were due to differences in fungal control, we quantified fungal burdens in target organs including kidney, liver and gastrointestinal tract at different time points. Unexpectedly, fungal burdens were comparable between KLRI1 or KLRI2-deficient mice and their WT controls across most tissues and time points examined (**Fig. 5d-e** and **Suppl. Fig. 5a-d**). The one exception was a modest but reproducibly significant increase in fungal burden in the kidneys of KLRI1-deficient mice at day 3 and 6 (**Fig. 5d** and **Suppl. Fig. 5c**). These results demonstrate that KLRI1 and KLRI2 exert opposing effects on host susceptibility to systemic *C. albicans* infection that is unrelated to major differences in fungal clearance from tissues.

### Differential renal immunopathology underlies opposing outcomes in KLRI1- and KLRI2-deficient mice

We next explored if there were alterations in tissue immunopathology that would explain the differential mortality observed in KLRI1- and KLRI2-deficient mice. We focused on the kidney, which is a primary target organ during systemic *C. albicans* infection, and mortality in murine models typically results from progressive renal failure caused by fungal proliferation and tissue destruction^17^. Overall, the total cellular infiltration and cytokines levels in the kidney were largely comparable in both knockout animals to those in WT controls (**Suppl. Fig. 6a-d**). The notable exceptions were a reduction in macrophages and increased IL-10 levels in the kidneys of the KLRI1-deficient mice (**Suppl. Fig. 6a-d**). There were no major differences in neutrophil activation and migration markers, except for a reduction in CD11b expression in KLRI2-deficient compared to WT mice (**Suppl. Fig. 6e-f**). During systemic *C. albicans* infection, renal macrophages are essential for fungal clearance^18^, but can also act as key drivers of kidney inflammation and tissue damage^19^. In this context, the reduction in macrophage numbers observed in KLRI1-deficient mice could account for the increased fungal burden in the kidney, while the concurrent rise in IL-10 levels may help limit excessive inflammation and tissue injury, thereby providing protection against immunopathology.

In terms of pathology, there were no detectable differences in the kidney between KLRI1-deficient and WT animals. In contrast, kidneys from KLRI2-deficient mice exhibited pronounced pathological alterations at day 9, characterised by increased inflammatory cell infiltration and associated tissue damage compared with WT mice (white arrows, **Fig. 6a**). Consistent with this, KLRI2-deficient mice displayed increased serum levels of the sepsis marker NGAL-2 at day 6 (**Fig. 6b**), pointing to heightened systemic inflammation. Moreover, KLRI2-deficient mice showed significantly elevated serum creatinine at day 9 post-infection, indicative of renal dysfunction (**Fig. 6c**). These markers were unaltered in the KLRI1-deficient mice. Together, these findings indicate that KLRI2 deficiency leads to exacerbated renal inflammation and tissue damage associated with systemic inflammation and kidney dysfunction, whereas KLRI1 deficiency preserves tissue integrity despite similar fungal burdens, highlighting distinct immunopathological mechanisms driving their opposing susceptibilities to systemic candidiasis.

**Fig. 6.**
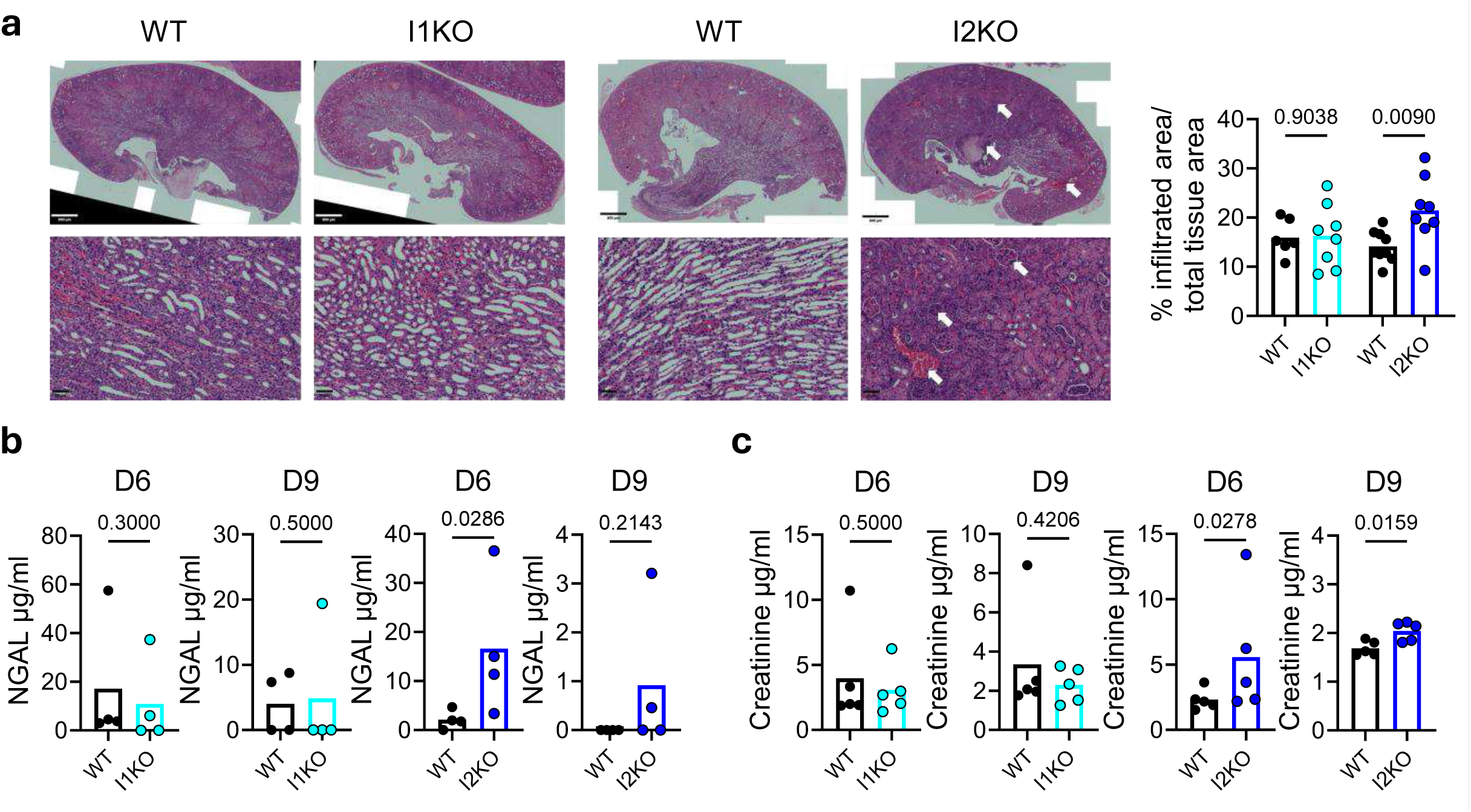
Differential renal immunopathology underlies opposing outcomes in KLRI1- and KLRI2-deficient mice. (a) H&E staining of formalin-fixed, paraffin-embedded kidney sections at day 9 post systemic *C. albicans* infection. Images shown are representative of two independent experiments. White arrows indicate areas of inflammatory cell infiltration and associated tissue damage. Quantification of the infiltrated area was performed using QuPath and expressed as the percentage of total kidney tissue area. Each point represents an individual mouse; data are presented as mean. **(b)** Serum NGAL (lipocalin-2) levels in WT, KLRI1- and KLRI2-deficient mice at days 6 and 9 post systemic *C. albicans* infection. Each point represents an individual mouse; bars indicate mean. Data are from one experiment. **(c)** Serum creatinine levels in WT, KLRI1- and KLRI2-deficient mice at days 6 and 9 post systemic *C. albicans* infection. Each point represents an individual mouse; bars indicate mean. Data are from one experiment. Statistical analysis was performed using Student’s t-test for two comparisons; *p < 0.05.

### Distinct systemic and T cell-mediated immune responses underlie opposing susceptibilities in KLRI1- and KLRI2-deficient mice

We then investigated whether there were alterations in systemic inflammatory responses by measuring serum cytokine profiles at days 6 and 9 post-*C. albicans* infection. At day 6, KLRI1-deficient mice showed significantly elevated IL-6 and reduced GM-CSF levels compared to WT controls (**Suppl. Fig. 7a**). The reduction in GM-CSF may account for the lower macrophage numbers in the kidneys of these mice, which could in turn contribute to the elevated fungal burden observed in this tissue. Although IL-6 typically promotes Th17 differentiation during systemic candidiasis^20,21^, we did not observe a corresponding increase in IL-17 production. A transient rise in IFN-y was also noted in KLRI1-deficient mice at day 6 but was not sustained (**Suppl. Fig. 7a**). In KLRI2-deficient mice, IL-4 levels were consistently elevated at both days 6 and 9, accompanied by increased IL-10 at day 6 and higher eotaxin (CCL11) at day 9 (**Suppl. Fig. 7a-b**), indicative of a Th2-skewed cytokine environment likely to impair antifungal immunity in the longer term.

To determine whether differential T cell responses contribute to the differential inflammatory response of the KLRI1- and KLRI2-deficient mice, we performed *in vitro* recall assays with splenocytes at day 9 and assessed intracellular cytokine production. Using *C. albicans* antigens to stimulate antigen-specific responses, we found no differences in T cell activation in KLRI1-deficient mice, whereas KLRI2-deficient mice displayed markedly enhanced T cell activation compared with WT, as indicated by a significant increase in CD44 expression (**Fig. 7a-b and Suppl. Fig. 7c**). Notably, KLRI2-deficient mice showed pronounced alterations in T cell polarization profiles, with a marked upregulation of IFN-y^+^, IL-4+ and IL-10+ CD4+ T cells (**Fig. 7c-d**). There were no major alterations in the T cell polarisation in the KLRI1-deficient mice, other than a reduction in IL-17A+ CD4+ T cells (**Fig. 7c-d**). Interestingly, we observed a trend toward an increased frequency of IL-10+ CD4+ T cells, which correlates with the elevated IL-10 levels in kidney homogenates, suggesting a more regulated immune response. Together, these findings indicate that despite having higher fungal burdens during infection, KLRI1-deficient mice mount a more balanced immune response, effectively limiting inflammation and tissue damage and thereby enhancing protection against infection. In contrast, loss of KLRI2 profoundly disrupts T cell and inflammatory responses, leading to excessive tissue damage and increased susceptibility to infection.

**Fig. 7.**
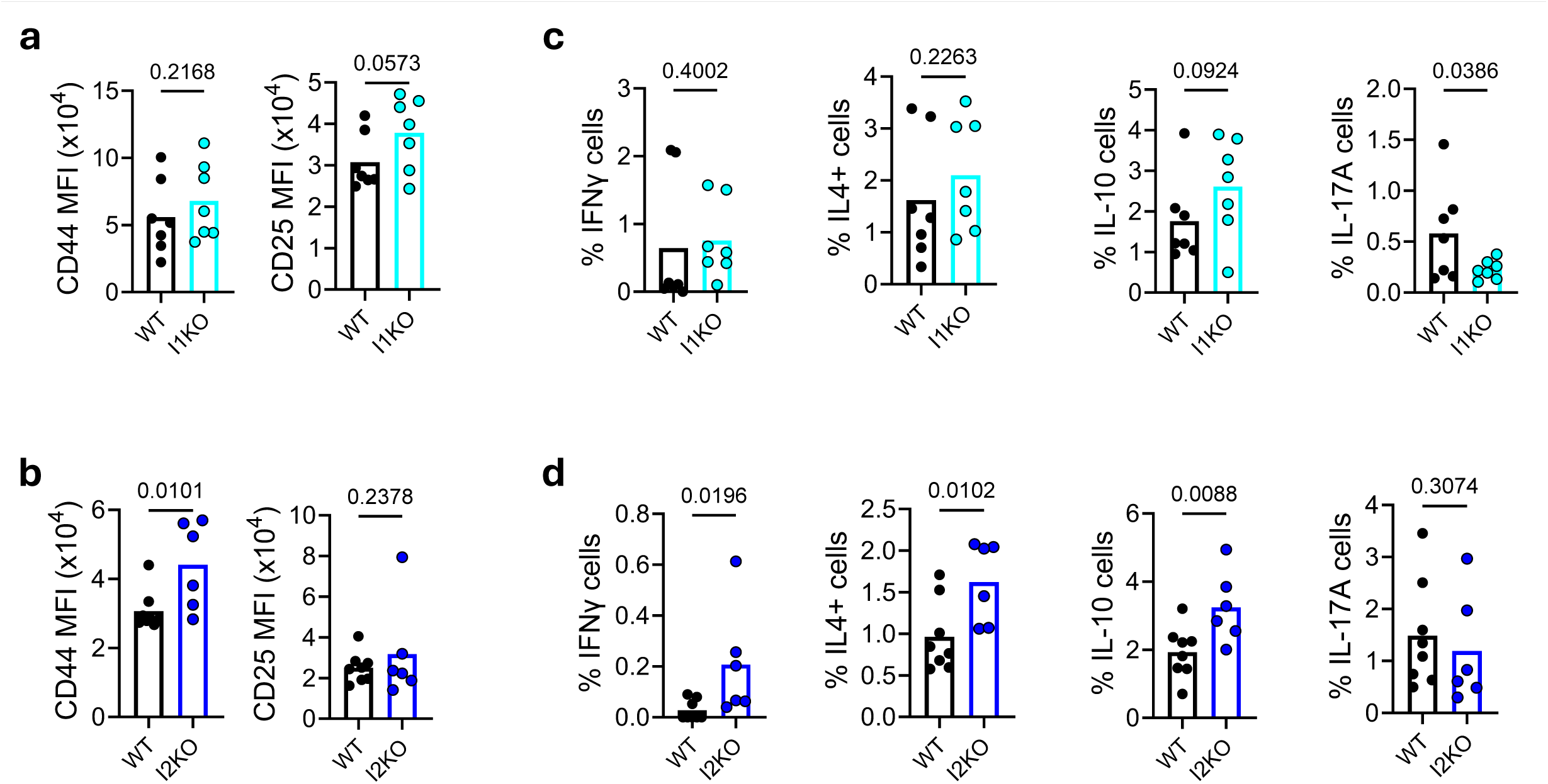
Distinct systemic and T cell-mediated immune responses underlie opposing susceptibilities in KLRI1- and KLRI2-deficient mice. (a) *In vitro* recall assay with HKCA stimulation of splenocytes from WT and KLRI1-deficient mice at day 9 post-infection. T cell activation (CD44, CD25) was assessed by flow cytometry. **(b)** *In vitro* recall assay with HKCA stimulation of splenocytes from WT and KLRI2-deficient mice at day 9 post-infection. T cell activation (CD44, CD25) was assessed by flow cytometry. **(c)** *In vitro* recall assay with HKCA stimulation of splenocytes from WT and KLRI1-deficient mice at day 9 post-infection. T cell intracellular cytokine production (IL-4, IL-10, IFN-y, IL-17A) was assessed by flow cytometry. **(d)** *In vitro* recall assay with HKCA stimulation of splenocytes from WT and KLRI2-deficient mice at day 9 post-infection. T cell intracellular cytokine production (IL-4, IL-10, IFN-y, IL-17A) was assessed by flow cytometry. Each point representsan individual mouse; bars indicate mean. Data are from one experiment. Statistical analysis was performed using Student’s t-test for two comparisons; *p < 0.05.

## Discussion

In this study, we identify KLRE1 as a novel and context-dependent modulator of DC activation and T cell priming, operating through distinct heterodimeric interactions with KLRI1 and KLRI2. While KLR family members have been extensively studied in NK cells, where they balance activating and inhibitory signals to fine-tune cytotoxicity and cytokine production^9^, their roles in DC biology have remained largely unexplored. Our data extend the functional reach of this receptor family to DCs, revealing a new layer of immune regulation at the intersection of innate recognition and adaptive T cell responses.

We show that KLRE1 is expressed by both tissue-resident and bone marrow-derived DCs, with expression dynamically regulated by the inflammatory environment. Notably, we find that TLR3 ligands, type I and II IFNs, and fungal stimuli such as HKCA strongly induce KLRE1 expression, suggesting that KLRE1 is positioned to respond to specific danger signals and to shape DC activation accordingly. This adds KLRE1 to a growing list of non-classical innate receptors whose expression is integrated with canonical PRR signalling pathways to fine-tune DC function^22^. Mechanistically, our data reveal that KLRE1 exerts bidirectional effects on DC activation through its distinct heterodimeric partners, KLRI1 and KLRI2. KLRE1 capping promotes the expression of key co-signalling molecules, including CD40, CD80, CD86, and MHC-II, in DCs. This effect is dependent on KLRI1, as KLRE1 capping in KLRI1-deficient DCs results in a reduced upregulation of these molecules compared with WT DCs. In contrast, both KLRE1 and KLRI2 deficiency independently result in elevated basal expression of these molecules, particularly CD80 and CD86, suggesting that the KLRE1:KLRI2 complex acts as a negative regulator of DC activation at steady state. This dual role mirrors the well-established paradigm in NK cells, where KLR heterodimers balance activating and inhibitory cues to control cytotoxic responses^9^. Here, we show that a similar regulatory logic extends to DCs, but with consequences for antigen presentation and T cell activation. One striking finding is that KLRI1 and KLRI2 are differentially regulated by specific inflammatory cues, shaping the availability of each heterodimer under different conditions. For example, TLR3 and type I IFNs upregulate both receptors, while IFN-γ selectively enhances KLRI1 expression. This suggests that during infection or inflammation, the local microenvironment could dynamically shift the balance of KLRE1 heterodimers, enabling context-specific modulation of DC activation.

KLRE1 does not contain any known signalling motifs and therefore requires heterodimerization with either KLRI1 or KLRI2 to transduce intracellular signals and regulate cellular functions^10,23^. KLRI1 has been characterised as an inhibitory receptor, containing immunoreceptor tyrosine-based inhibitory motifs (ITIMs) in its cytoplasmic domain^10,23^, while KLRI2 lacks these motifs and is considered an activating receptor, likely signalling through adaptor proteins bearing ITAMs^10,23^. However, our study reveals that the KLRE1:KLRI1 heterodimer promotes DC activation, while KLRE1:KLRI2 heterodimers act as a negative regulator. One possibility to explain our findings is that the KLRE1:KLRI1 and KLRE1:KLRI2 heterodimers might both ultimately act on a shared inhibitory pathway within DCs, but with opposing effects on its activity. In this model, KLRI1 could function by recruiting phosphatases or other inhibitory mediators that suppress a dominant intracellular inhibitor of DC activation. Thus, the KLRE1:KLRI1 complex would paradoxically promote DC activation by inhibiting an inhibitor, resulting in a net positive effect on co-signalling molecule expression and T cell priming. Conversely, KLRI2, traditionally classified as an activating receptor, may instead amplify the activity of the same or another intracellular inhibitory pathway when paired with KLRE1 in DCs. In this scenario, the KLRE1:KLRI2 heterodimer would reinforce a negative regulatory circuit, dampening co-stimulatory molecule expression under steady-state conditions and acting as a brake on DC activation. Such an arrangement is conceptually similar to regulatory circuits in other immune pathways, where ITIM-bearing receptors can paradoxically promote activation by dampening dominant negative feedback loops^24-26^. For example, in some settings, inhibitory receptors such as PD-1 and CTLA-4 act by constraining key negative regulators of TCR signalling^24,25^, and certain Fcy receptors can switch between activating and inhibitory functions depending on adaptor availability and ligand context^26^. Our findings suggest that KLRE1 heterodimers may operate under a similar logic in DCs, with their net outcome on cell activation determined not only by receptor motifs but by which intracellular pathways they modulate and how these pathways are wired into the broader signalling network. Future studies dissecting the downstream signalling partners of KLRE1:KLRI1 and KLRE1:KLRI2 in DCs, particularly their interactions with known inhibitory phosphatases, adaptors, or transcriptional repressors, will be crucial to test this hypothesis.

Functionally, we show that KLRE1 deficiency enhances DC activation and boosts T cell responses both *in vitro* and *in vivo*. Interestingly, this increased activation does not translate into improved pathogen clearance during systemic *Candida albicans* infection. Instead, our data point to a decoupling of immune activation from fungal burden in this setting, highlighting a potential role for KLRE1 in tuning not just resistance but also disease tolerance. Supporting this, KLRI1- and KLRI2-deficient mice show inverse survival outcomes during systemic candidiasis with a moderate but significant increase in fungal load in the kidneys of KLRI1-deficient mice. Notably, KLRI2-deficient mice exhibited systemic inflammation, elevated sepsis markers, and renal dysfunction, indicating that their increased susceptibility to systemic *C. albicans* infection is largely driven by pathologic inflammatory response. Increased CD44 expression and elevated cytokine production in T cells indicate that their heightened activation may exacerbate pathology, weakening protective responses while amplifying tissue damage. In contrast, KLRI1 deficiency was associated with a more controlled and regulated T cell response, characterised by elevated IL-10 levels in kidney homogenates and a reduction in macrophage numbers. This combination may help limit excessive inflammation and tissue damage, thereby promoting survival despite higher fungal burdens in the tissues. Together, these findings highlight the role of KLRE1 heterodimers in maintaining the delicate balance between protective immunity and immunopathology, with KLRI1 promoting disease tolerance through regulated T cell activation, whereas KLRI2 deficiency drives systemic inflammation and kidney damage through T cell hyperactivation.

Collectively, these findings expand our understanding of how non-classical innate receptors fine-tune DC function, acting as context-dependent checkpoints that integrate pathogen- and cytokine-derived signals to shape co-signalling, T cell polarisation, and ultimately host outcome. Our work aligns with a broader concept that immune homeostasis during infection depends not only on pathogen elimination (resistance) but also on the host’s ability to limit collateral damage (tolerance). KLRE1 and its heterodimeric partners appear to operate precisely at this interface. These results raise several important questions for future investigation. It will be essential to identify the molecular ligands on T cells or other interacting cells that engage KLRE1 heterodimers in DCs and to map the downstream signalling pathways associated with KLRE1:KLRI1 and KLRE1:KLRI2 complexes that mediate their respective activating and inhibitory effects. Furthermore, given that these receptors are also expressed on other immune cell types, such as lymphocytes, future studies using myeloid- or DC-specific knockout models will be critical to dissect the cell type-specific roles of these receptor pairs in regulating immune responses. Additionally, whether KLRE1 modulates DC-driven T cell responses in other contexts, such as cancer, autoimmunity, or vaccination, remains an open question with significant translational implications. Finally, our finding that manipulating KLRE1 engagement can boost or dampen co-stimulatory molecule expression suggests that targeting this pathway could provide new strategies to enhance DC-based immunotherapies or limit harmful inflammation.

In summary, our study establishes KLRE1 as a novel, dual-function modulator of DC phenotype and T cell activation, acting through distinct heterodimeric interactions to balance immunity and tolerance during fungal infection. These insights extend the immunological roles of KLRs beyond NK cells and uncover new avenues for modulating DC function to shape protective or tolerogenic immune responses.

## Methods

### Mice

Wild-type C57BL/6J, KLRE1-/- (provided by the RIKEN BRC through the National BioResource Project of the MEXT, Japan^11^), KLRI1-/- (and KLRI2-/- mice were maintained under specific pathogen-free (SPF) conditions at Charles River Laboratories. KLRI1-/- mice were generated as described before (co-submitted manuscript^8^). KLRI2 full knockout mouse model (C57BL/6N background) was generated using CRISPR/Cas9-mediated genome engineering (Cyagen, USA). Briefly, mouse genomic fragments containing homology arms were amplified from a BAC clone using high-fidelity Taq DNA polymerase and sequentially assembled into a targeting vector together with recombination sites and selection markers. Exon 1 was selected as the conditional knockout region. Guide RNA (gRNA) targeting the Klri1 gene and Cas9 mRNA were co-injected into fertilized mouse zygotes to generate knockout offspring. F0 founder mice were identified by PCR and confirmed by sequence analysis. The line was subsequently rederived onto a C57BL/6N background at Charles River Laboratories. Hector TCR-transgenic and OT-II TCR-transgenic mice were bred at the University of Exeter and housed under SPF conditions. Animals were kept in individually ventilated cages with food and water ad libitum, on a 12 h light/dark cycle (20–24 °C, 50–60 % humidity). Mice used in experiments were 6–8-week-old females, co-housed for ≥14 days prior to experiments and randomly assigned to groups. All procedures were conducted under UK Home Office licence and approved by the University of Exeter Animal Welfare and Ethical Review Body (project license number: PP9965358).

### Systemic *Candida albicans* infection

Mice were injected intravenously with 1 × 10⁵ CFU *C. albicans* SC5314, with or without 1 × 10⁶ Hector CD4+ T cells (transferred 24 h prior to infection). Yeasts were grown in YPD broth at 30 °C for 24 h, washed twice in PBS, counted, and adjusted to the required concentration. Kidneys, gastrointestinal tract and liver were collected at the indicated time points, homogenised in PBS, serially diluted, and plated on YPD agar supplemented with gentamicin (100 µg/ml) and vancomycin (10 µg/ml). Plates were incubated for 24 h at 37 °C before CFU enumeration. Clinical status and weight were monitored daily; humane endpoints were applied for >30% weight loss or severe illness.

### Tissue digestion and single-cell suspension

mLN and spleens were gently dissociated through 100 µm strainers into RPMI-1640 with 10% Foetal Calf Serum (FCS). Spleen red blood cells were lysed with 1x RBC lysis buffer (BD PharmLyse, BD Biosciences). Bone marrow was collected by flushing femurs and tibiae with RPMI-1640 and filtering through 100 µm strainers. Kidneys were minced with scissors, digested for 30 min at 37 °C with Liberase TL (Roche, 1 mg/ml) and DNase I (Merck, 50 µg/ml), and filtered. All suspensions were washed (400 *×g*, 5 min, 4 °C) and resuspended in complete RPMI (10% FCS, 1% penicillin/streptomycin, 50µM 2-mercaptoethanol) for tissue culture, or FACS buffer (PBS with 10% FCS, 2mM EDTA) for flow cytometry analysis.

### Bone marrow-derived dendritic cells (BMDCs) GM-CSF/IL-4 cultures

Bone marrow cells were plated at 1 × 10⁶ per ml in DMEM-GlutaMAX with 10% FCS, 1% penicillin/streptomycin, 50 µM 2-mercaptoethanol and GM-CSF (20 ng/ml; Biolegend) with or without IL-4 (10 ng/ml; Bio-techne). Fresh medium was added on day 3, and cells were harvested on day 7.

### Flt3L/OP9 cultures

Bone marrow cells were seeded at 4 × 10⁶ cells/ml in RPMI-1640 supplemented with 10% FCS, 1% penicillin/streptomycin, 50 µM 2-mercaptoethanol, and Flt3L (200 ng/ml; BioLegend). Cultures were maintained for 7 days, with fresh medium added on days 3–4. For OP9 co-cultures, cells were transferred on days 3–4 onto a confluent monolayer of mitomycin C–treated (Stell Cell Technologies) OP9 stromal cells engineered to express the Notch ligand Delta-like 1 (DLL1) (kindly donated by Hans R. Haecker, University of Utah). Non-adherent cells were harvested on day 7 for analysis. OP9 cells were maintained separately in α-MEM supplemented with 20% FCS and 1% penicillin/streptomycin.

### BMDC stimulation

BMDCs generated from the above culture methods were plated at 2,5 × 10^5^ per ml and stimulated with poly(I:C) (1 µg/ml), CpG ODN (1 µg/ml), LPS (1 µg/ml), FSL-1 (0.5 µg/ml), Pam₂CSK₄ (1 µg/ml) (all from InvivoGen), recombinant murine IFN-α, IFN-β, IFN-γ or IFN-λ (100 ng/ml, Biolegend), heat-killed *C. albicans* SC5314 (MOI 1:1), furfurman (10 µg/ml) or curdlan (10 µg/ml; (all from InvivoGen). In some experiments, BMDCs were stimulated with plate-bound rat anti-mouse KLRE1 antibody. Briefly, plates were first coated with anti-rat IgG antibody (1 µg/ml; Thermo Fisher Scientific) for 1 h at 37 °C, followed by rat anti-mouse KLRE1 antibody (1 µg/ml; Bio-Techne) for an additional 1 h at 37 °C. After stimulation, cells were harvested at 16 h for RNA extraction or at 24 h for cytokine measurement and flow cytometry analysis.

### Flow cytometry

Single-cell suspensions were prepared in FACS buffer and stained with fixable viability dye (eFluor 780; eBioscience), washed, then stained with fluorochrome-conjugated monoclonal antibodies for same-day acquisition or fixed with 2 % PFA (Sigma). Conjugated monoclonal antibodies included: CD80-FITC/BV650 (16-10A1, BioLegend), CD86-AF700 (GL1, eBioscience), CD40-BV605 (3/23, BioLegend), MHC-II-BUV496 (2G9, BD Biosciences), CD44-FITC/BV510 (IM7, BD Biosciences/BioLegend), CD25-BV650/PE-Cy7 (PC61, BD Biosciences/BioLegend), CD90.1-BB700/APC (HIS51, OX-7; BD Biosciences), CD45-BUV496/PerCP-Cy5.5 (30-F11, BD Biosciences/BioLegend), CD4-BUV563/BV510 (RM4-5, RM4-4; BD Biosciences/BioLegend), CD3-AF594/eFluor 450 (500A2, BioLegend/eBioscience), B220-eFluor 450/AF700 (RA3-6B2, eBioscience/BioLegend), CD49b-eFluor 450 (DX5, eBioscience), CD11b-BUV395 (M1/70, BD Biosciences), Ly6G-eFluor450/SB550 (1A8, eBioscience/BioLegend), Ly6C-BV570 (HK1.4, BioLegend), F4/80-FITC/BV421 (BM8, BioLegend), CD11c-BV711 (HL3, BD Biosciences), NKp46-SB645 (29A1.4, eBioscience), NK1.1-BV605/AF647 (PK136, HP-3G10; BD Biosciences/BioLegend), Siglec-F-SB436 (1RNM44N, eBioscience), CD62L-BV605/PE/BUV563 (MEL-14, BD Biosciences), CCR1-PE (643854, R&D Systems), CXCR2-AF647 (SA045E1, BioLegend), CD18-APC (H155-78, BioLegend), CX3CR1-BV785/PE (SA011F11, BioLegend), KLRE1-AF488 (854929, Bio-techne), PDCA-1-BUV563 (927, BD Biosciences), CD8-AF532 (RPA-T8, Thermo Fisher Scientific), CD64-AF647 (X54-5/7.1, BD Biosciences) and CD26-BUV737 (H194-112, BD Biosciences). For intracellular staining, T-bet-BV421/BV786 (O4-46, BD Biosciences), GATA-3-PE-Cy7/BUV395 (L50-853, BD Biosciences), FoxP3-AF488/AF647 (MF23, BD Biosciences), RORyt-BV650/PE (Q31-378, BD Biosciences), IFN-y-FITC (XMG1.2, BD Biosciences), IL-4-BV605 (11B11, BD Biosciences), IL-10-PE (JES5-16E3, BD Biosciences) and IL-17A- AF700 (TC11-18H10, BD Biosciences) were used. Intracellular staining employed the Transcription Factor Staining Buffer Set as per manufacturers instructions (eBioscience). Data were acquired on a Attune NxT (Thermo Fisher Scientific) or Cytek Aurora and analysed with FlowJo v10 (BD Biosciences).

### *In vitro* T cell–dendritic cell co-culture

CD11c+ DCs were harvested from BMDCs cultures or purified from mLN of wild-type or Klre1-/- mice using CD11c MicroBeads UltraPure (Miltenyi Biotec), as per manufacturers instructions. OT-II CD4+ T cells were purified from spleens and lymph nodes be negative selection using the EasySep Mouse CD4^+^ T Cell Isolation Kit (STEMCELL Technologies), as per manufacturers instructions. DCs were pre-incubated with ovalbumin (OVA) peptide (2 µg/ml, InvivoGen) for 1 h before co-culture with T cells at a 5:1 ratio. Where indicated, KLRE1 antibody (10 µg/ml, Bio-techne) or isotype control was added. After 48–96 h, cells were harvested an analysed by flow cytometry.

### Histology

Kidneys were fixed in 10% neutral-buffered formalin, paraffin-embedded, sectioned at 4 µm, and stained with Hematoxylin and Eosin then counterstained with Periodic acid–Schiff (PAS). Processing was performed by Microtechnical Services Ltd. (Exeter, UK). Images were taken using a APEXVIEW APX100 fluorescence microscope. Quantification of the infiltrated area was performed using QuPath.

### Quantitative PCR

RNA was extracted from mLN, spleen and BMDCs using TRIzol (Thermo Fisher Scientific), as per manufacturers instructions, quantified on a NanoDrop (Thermo Fisher Scientific), and reverse-transcribed with SuperScript III into cDNA (Thermo Fisher Scientific), as per manufacturers instructions.

SYBR Green assays used PowerTrack SYBR Green Master Mix (Thermo Fisher Scientific) as per manufacturers instructions on a QuantStudio 7 Pro (Applied Biosystems). Cycling was 95 °C for 2 min, followed by 40 cycles of 95 °C for 5 s and 60 °C for 30 s, with melt-curve analysis. Primer sequences were: Klre1 (F 5′-GTCAGTTTGTTCGCACAG-3′, R 5′-TGAAACTCGCAGACAGTG-3′), Klri1 (F 5′-GGTAACTGGCATGCTTGGAG-3′, R 5′-CAAAGATCACAGGAGGCGTC-3′), Klri2 (F 5′-ACCCTCGAAGACAGACCAGAG-3′, R 5′-CCACATCGAATCCTGTCCTG-3′), Cd36 (F 5′-GTCCAAGTCTTCTATGTTCC-3′, R 5′-CTCCATCTACAGTGTCATTG-3′), Il12b (F 5′-TTCAACATCAAGAGCAGTAG-3′, R 5′-AGGACACTGAATACTTCTCA-3′), Relb (F 5′-CTTCTCTCAAGCTGATGTG-3′, R 5′- AAGAACACATTGACAGTCAC-3′). Reference genes: Gapdh (F 5′-ACGACCCCTTCATTGACCTC-3′, R 5′-CATTCTCGGCCTTGACTGTG-3′) and 18srrna (F 5′-ATGAACGAGGAATTCCCAG-3′, R 5′-CCAATCGGTAGTAGCGAC-3′). Expression was normalised to Gapdh and 18srrna and analysed by the ΔΔCt method.

In addition, the Mouse Dendritic Cell TaqMan® Array 96 Well Plate (Applied Biosystems) were used. Assays were run with TaqMan Universal PCR Master Mix as per manufacturers instructions.

### ELISA

Murine blood was collected via cardiac puncture, clotted, centrifuged (10,000 × g, 10 min), and serum was stored at –80 °C. Creatinine (abcam) and lipocalin-2 (R&D Systems) were measured with mouse ELISA kits as per manufacturers instructions. IL-2 from supernatants from T cell–DC co-cultures were quantified using an IL-2 DuoSet ELISA (R&D Systems) as per manufacturers instructions. Absorbance was read at 450 nm with 570 nm correction on a Spark microplate reader (TECAN).

### Luminex multiplex cytokine analysis

Serum cytokines (IL-1β, TNF, IFN-γ, IL-2, IL-6, IL-4, MCP-1, IL-5, IL-10, KC, IL-17A, GM-CSF, eotaxin) were measured using Bio-Plex Pro Mouse Cytokine kits (Bio-Rad) as per manufacturers instructions. Plates were read on a Luminex MAGPIX instrument and analysed with Bio-Plex Manager software.

### *In vitro* T cell restimulation

Splenocytes (2 × 10⁶ per well) were cultured in 96-well plates. For polyclonal stimulation, cells were treated with PMA (Sigma, 50 ng/ml) and ionomycin (500 ng/ml; Sigma) for 3 h, followed by brefeldin A (10 µg/ml, BioLegend) for 3 h. For fungal restimulation, splenocytes were incubated with heat-killed *C. albicans* SC5314 (MOI 1:1) for 72 h. Cells were analysed by intracellular cytokine staining and flow cytometry.

### Statistical analysis

Statistical analyses were performed using GraphPad Prism 10. Fungal burden data were evaluated with Mann–Whitney U tests. Survival data were analysed with Kaplan-Meier test. For single comparisons involving one variable Student’s t-tests were used, while One-way ANOVA was applied for analyses involving multiple groups with Dunnett’s multiple comparison test. Two-way ANOVA was employed when assessing the effect of two independent variables.

Data are presented as mean unless otherwise specified. A P-value < 0.05 was considered statistically significant.

## Acknowledgement

We gratefully acknowledge the Exeter Centre for Cytomics for their support with flow cytometry analysis and the University of Exeter Biological Services Unit for their assistance with animal experiments. This work was supported by the MRC Centre for Medical Mycology at the University of Exeter (MR/N006364/2 and MR/V033417/1), the NIHR Exeter Biomedical Research Centre, and the Wellcome Trust (217163/Z/19/Z). The views expressed are those of the author(s) and not necessarily those of the NIHR or the Department of Health and Social Care. For the purpose of open access, the author has applied a CC BY public copyright licence to any Author Accepted Manuscript version arising from this submission.

---

**Suppl. Fig. 1.**
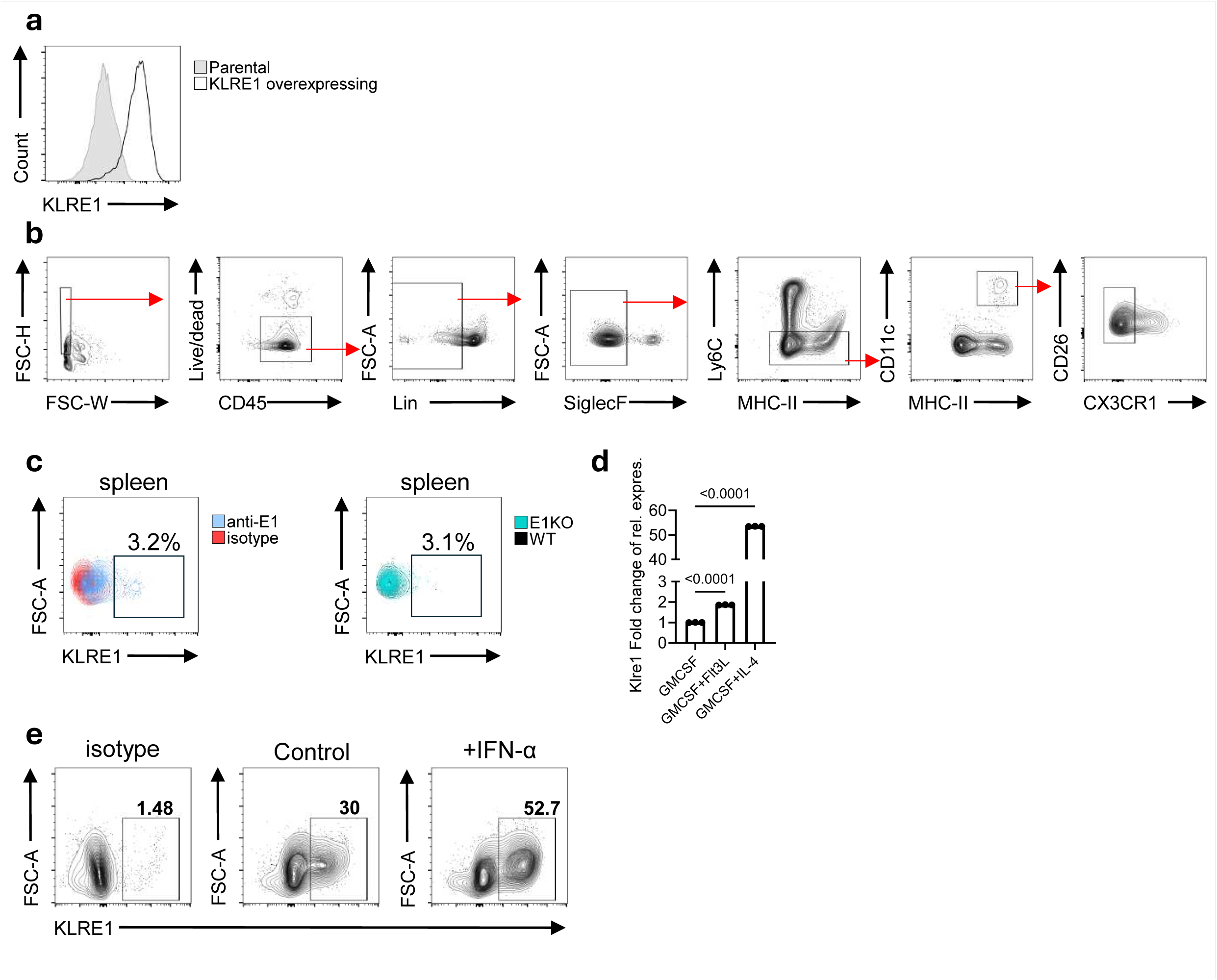
KLRE1 is expressed by DCs and regulated by inflammatory signals. **(a)** Flow cytometric analysis of KLRE1 expression in parental and KLRE1 overexpressing NIH3T3 cells. **(b)** Gating strategy used to identify KLRE1+ cells within tissue DCs. **(c)** Flow cytometric analysis of KLRE1 expression in spleen DCs (gated on viable CD45+ Lin- CD11c+ MHC-II+ CX3CR1-). KLRE1 expression was assessed with a specific anti-KLRE1 monoclonal antibody (blue), using an isotype control for background (red), and compared to expression in KLRE1-EGFP knock-in mice (blue) with WT mice as controls (red). **(d)** KLRE1 gene expression in bone marrow–derived cells cultured for at least 7 days with different cytokine combinations. Data are shown as expression relative to GM-CSF treated samples and normalized to housekeeping genes. Bars represent the mean from one experiment with triplicates, shown as representative of at least two independent experiments. **(e)** Flow cytometric analysis of KLRE1 expression in FD-BMDCs after 24 h stimulation with IFN-a. rLRE1 expression was assessed with a specific anti-KLRE1 monoclonal antibody, using an isotype control for background. Statistical analysis was performed using Student’s t-test for two comparisons and One-way ANOVA for three or more comparisons; *p < 0.05.

**Suppl. Fig. 2.**
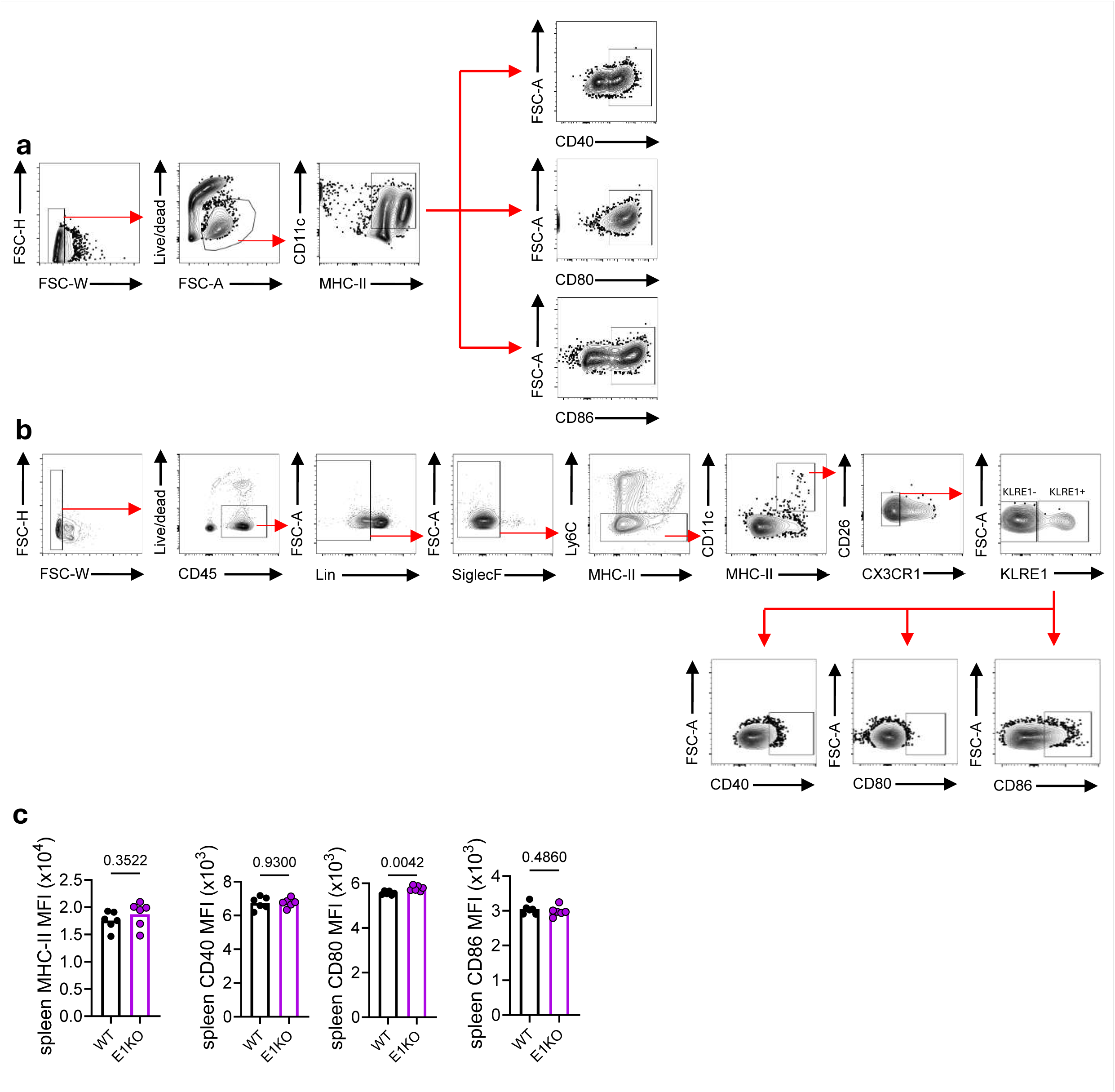
KLRE1 alters DC phenotype by regulating co-signalling molecule expression. **(a)** Gating strategy for assessing MHC-II and co-signalling molecule expression (CD40, CD80, CD86) in FD-BMDCs. **(b)** Gating strategy for assessing MHC-II and co-signalling molecule expression (CD40, CD80, CD86) in tissue DCs (mLN and spleen). **(c)** Flow cytometric analysis of MHC-II, CD40, CD80, and CD86 expression in spleen DCs at steady state. Each point represents one mouse; horizontal bars indicate the mean. Data show MFI of pooled data from at least two independent experiments. Statistical analysis was performed using Student’s t-test for two comparisons; *p < 0.05.

**Suppl. Fig. 3.**
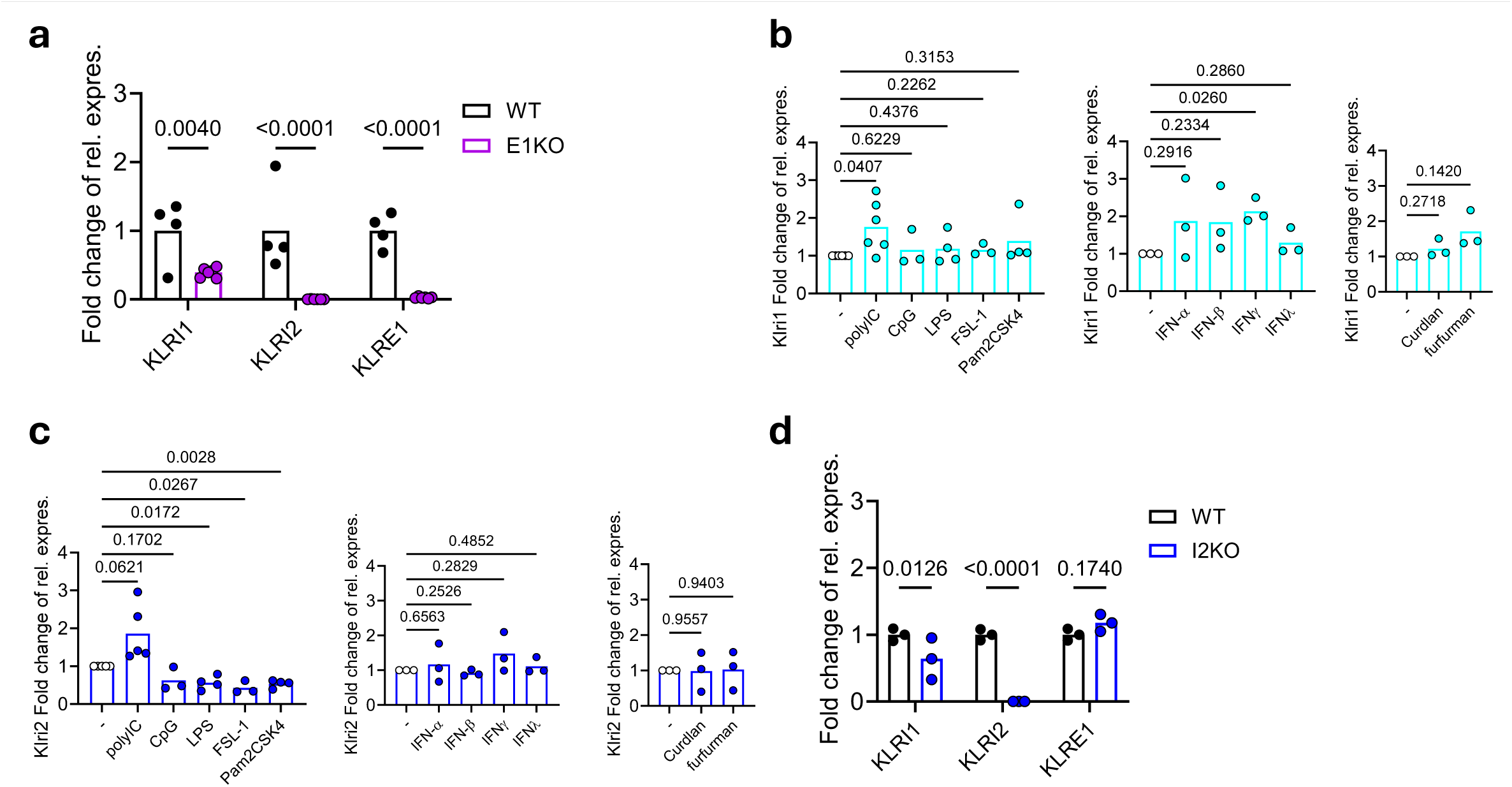
KLRE1 modulates DC activation via distinct heterodimeric interactions with KLRI1 and KLRI2. **(a)** KLRI1, KLRI2 and KLRE1 transcript levels in mLN of WT and KLRE1-deficient mice at steady state. Data are shown relative to WT controls and normalised to housekeeping genes. Bars represent the mean. **(b)** KLRI1 mRNA expression in FD-BMDCs after 16 h stimulation with the indicated TLR ligands, CLR agonists, or interferons. Data are shown relative to unstimulated controls and normalised to housekeeping genes. Bars represent the mean of pooled data from of at least three independent experiments. **(c)** KLRI2 mRNA expression in FD-BMDCs after 16 h stimulation with the indicated TLR ligands, CLR agonists, or interferons. Data are shown relative to unstimulated controls and normalised to housekeeping genes. Bars represent the mean of pooled data from of at least three independent experiments. **(d)** KLRI1, KLRI2 and KLRE1 transcript levels in mLN of WT and KLRI2-deficient mice at steady state. Data are shown relative to WT controls and normalised to housekeeping genes. Bars represent the mean. Statistical analysis was performed using Student’s t-test for two comparisons and One-way ANOVA for three or more comparisons; *p < 0.05.

**Suppl. Fig. 4.**
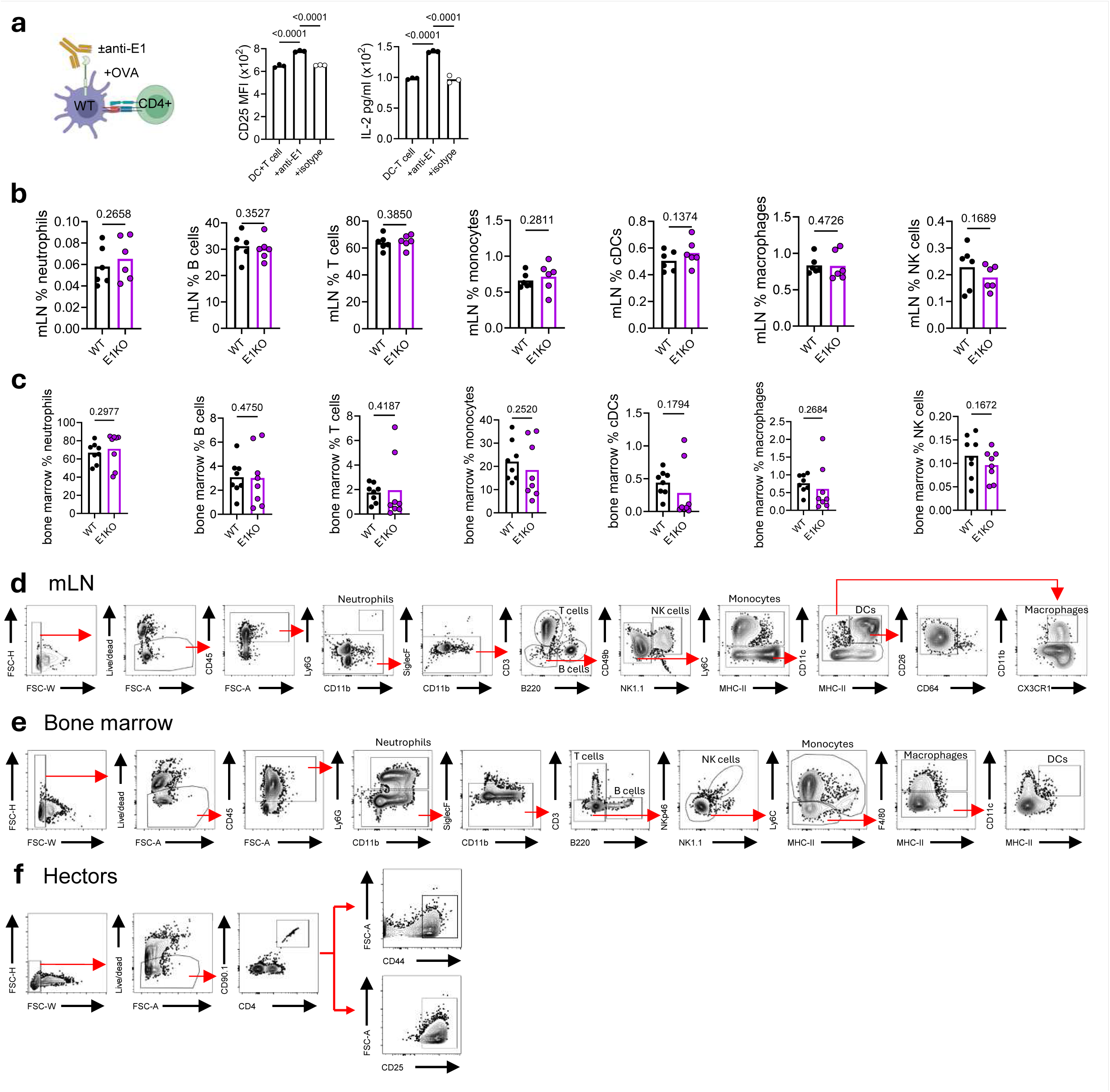
KLRE1 regulation of DC phenotype impacts T cell activation during fungal infection. **(a)** *In vitro* co-culture of WT DCs preloaded with OVA and OT-II CD4+ T cells for 72 h in the presence of anti-KLRE1 antibody or isotype control. T cell activation was assessed by CD25 expression and IL-2 production. Data show MFI from one experiment, shown as representative of at least two independent experiments. **(b)** Immune cell profiling of KLRE1-deficient mice in mLN. Each point represents an individual mouse; horizontal bars indicate the mean. Data are presented as the percentage of each cell population within total CD45+ viable cells. **(c)** Immune cell phenotyping of KLRE1-deficient mice in bone marrow. Each point represents an individual mouse; horizontal bars indicate the mean. Data are presented as the percentage of each cell population within total CD45+ viable cells. **(d)** Gating strategy used for immunophenotyping in mLN. **(e)** Gating strategy used for immunophenotyping in bone marrow. **(f)** Gating strategy for assessing Hector CD4+ T cell numbers, frequencies, and activation status (CD44 and CD25 expression) in the spleen following systemic *C. albicans* infection. Statistical analysis was performed using Student’s t-test for two comparisons; *p < 0.05.

**Suppl. Fig. 5.**
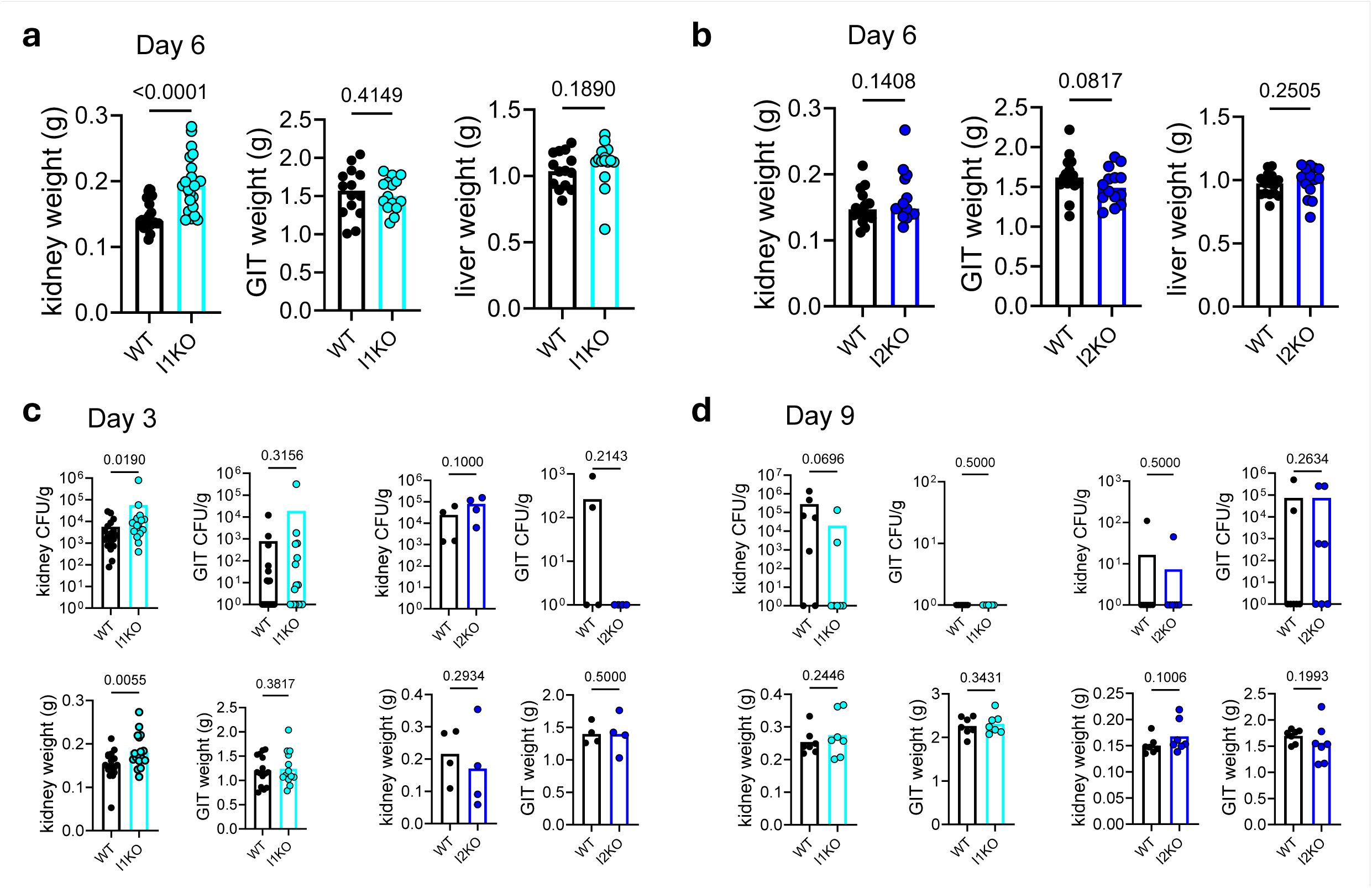
KLRI1 and KLRI2 inversely influence susceptibility to systemic *Candida* infection. **(a)** Tissue weights of kidney, GIT, and liver of WT and KLRI1-deficient mice at day 6 post-infection. Each point represents an individual mouse; bars indicate mean. Data pooled from at least two independent experiments, with ten mice per group in each experiment. **(b)** Tissue weights of kidney, GIT, and liver of WT and KLRI2-deficient mice at day 6 post-infection. Each point represents an individual mouse; bars indicate mean. Data pooled from two independent experiments, with ten mice per group in each experiment. **(c)** Fungal burden and tissue weights of kidney and GIT of WT, KLRI1 and KLRI2-deficient mice at day 3 post-infection. Each point represents an individual mouse; bars indicate meam. Data pooled from three (KLRI1 data) or one (KLRI2 data) independent experiments. **(d)** Fungal burden and tissue weights of kidney and GIT of WT, KLRI1 and KLRI2-deficient mice at day 9 post-infection. Each point representsan individual mouse; bars indicate mean. Data are from one experiment. Statistical analysis was performed using Student’s t-test for two comparisons. Fungal burden data were evaluated with Mann–Whitney U tests. *p < 0.05.

**Suppl. Fig. 6.**
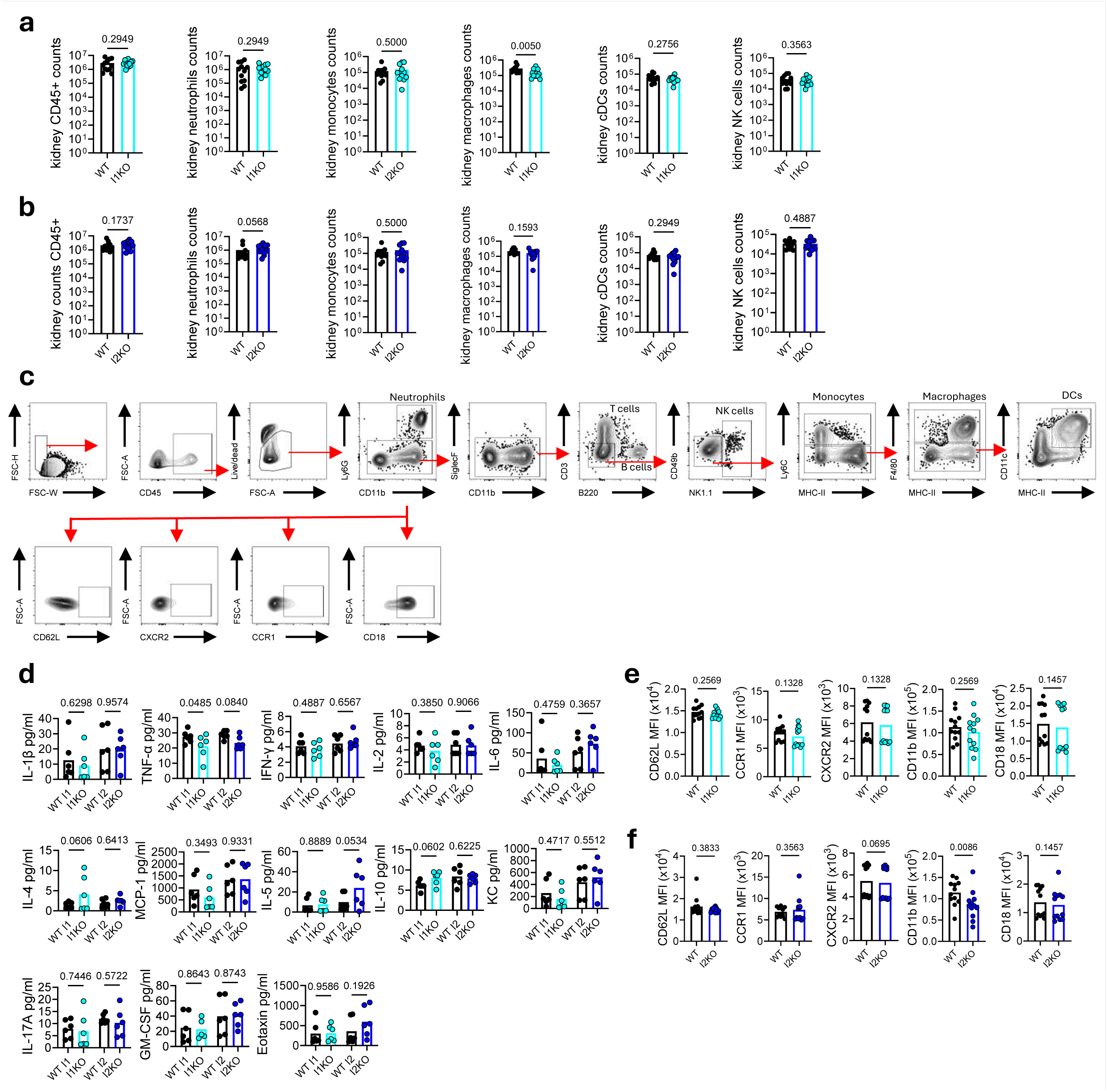
Differential renal immunopathology underlies opposing outcomes in KLRI1- and KLRI2-deficient mice. **(a)** Kidney innate immune cell recruitment in WT and KLRI1-deficient mice at day 6 post systemic *C. albicans* infection. Each point represents an individual mouse; bars indicate mean. Data show cell counts pooled from two independent experiments. **(b)** Kidney innate immune cell recruitment in WT and KLRI2-deficient mice at day 6 post systemic *C. albicans* infection. Each point represents an individual mouse; bars indicate mean. Data show cell counts pooled from two independent experiments. **(c)** Gating strategy for assessing innate immune cell recruitment and neutrophil activation in the kidneys following systemic *C. albicans* infection. **(d)** Cytokine production (IL-1̋, TNFa, IFN-y, IL-2, IL-4, IL-5, IL-6, IL-10, MCP-1 (CCL2), KC (CXCL1), IL-17A, GM-CSF and eotaxin (CCL11)) in kidney homogenates from WT, KLRI1- and KLRI2-deficient mice at day 9 post systemic *C. albicans* infection. Each point represents an individual mouse; bars indicate mean. Data are from one experiment. **(e)** Flow cytometric analysis of CD62L, CCR1, CXCR2, CD18 and CD11b expression in neutrophils from the kidneys from WT and KLRI1-deficient mice at day 6 post systemic *C. albicans* infection. Each point represents an individual mouse; bars indicate mean. Data show MFI pooled from two independent experiments. **(f)** Flow cytometric analysis of CD62L, CCR1, CXCR2, CD18 and CD11b expression in neutrophils from the kidneys from WT and KLRI2-deficient mice at day 6 post systemic *C. albicans* infection. Each point represents an individual mouse; bars indicate mean. Data show MFI pooled from two independent experiments. Statistical analysis was performed using Student’s t-test for two comparisons; *p < 0.05.

**Suppl. Fig. 7.**
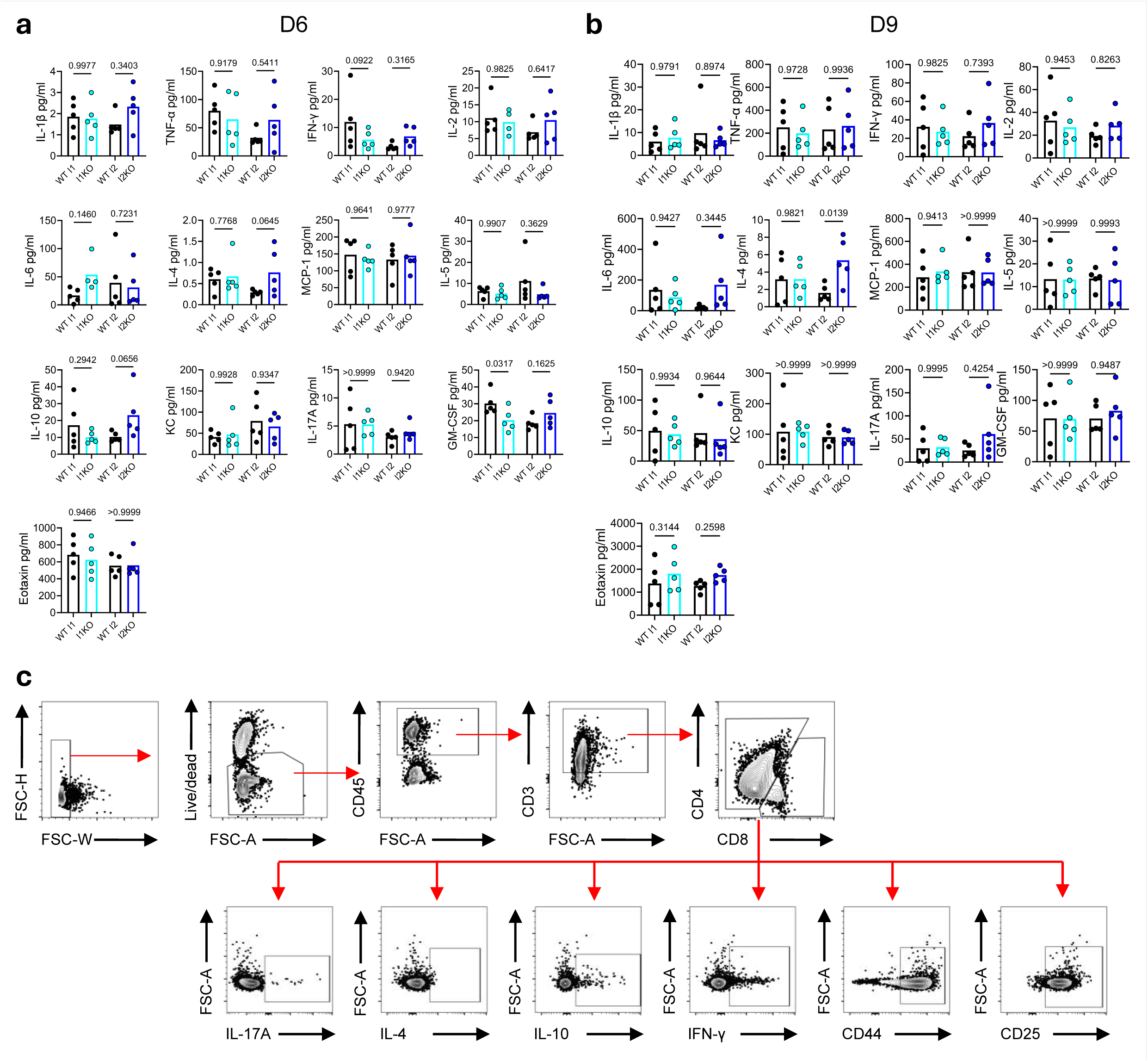
Distinct systemic and T cell-mediated immune responses underlie opposing susceptibilities in KLRI1- and KLRI2-deficient mice. **(a)** Cytokine (IL-1̋, TNFa, IFN-y, IL-2, IL-4, IL-5, IL-6, IL-10, MCP-1 (CCL2), KC (CXCL1), IL-17A, GM-CSF and eotaxin (CCL11)) in serum from WT, KLRI1- and KLRI2-deficient mice at day 6 post-infection. Each point represents an individual mouse; bars indicate mean. Data are from one experiment. **(b)** (a) Cytokine (IL-1̋, TNFa, IFN-y, IL-2, IL-4, IL-5, IL-6, IL-10, MCP-1 (CCL2), KC (CXCL1), IL-17A, GM-CSF and eotaxin (CCL11)) in serum from WT, KLRI1- and KLRI2-deficient mice at day 9 post-infection. Each point represents an individual mouse; bars indicate mean. Data are from one experiment. **(c)** Gating strategy for assessing T cell intracellular cytokine expression. Statistical analysis was performed using Student’s t-test for two comparisons; *p < 0.05.

## Notes

### Competing Interest Statement

The authors have declared no competing interest.

